# YTHDC2 Is Essential for Pachytene Progression and Prevents Aberrant Microtubule-Driven Telomere Clustering in Male Meiosis

**DOI:** 10.1101/2021.04.12.439470

**Authors:** Rong Liu, Seth D. Kasowitz, David Homolka, N. Adrian Leu, Jordan T. Shaked, Gordon Ruthel, Devanshi Jain, Scott Keeney, Mengcheng Luo, Ramesh S. Pillai, P. Jeremy Wang

**Affiliations:** School of Basic Medical Sciences, Wuhan University, Wuhan, Hubei Province, China; Department of Biomedical Sciences, University of Pennsylvania School of Veterinary Medicine, Philadelphia, Pennsylvania, USA; Department of Molecular Biology, Science III, University of Geneva, CH-1211 Geneva 4, Switzerland; Department of Pathobiology, University of Pennsylvania School of Veterinary Medicine, Philadelphia, Pennsylvania, USA; Molecular Biology Program, Memorial Sloan Kettering Cancer Center, New York, USA; Department of Genetics, Rutgers University, Piscataway, NJ, USA; Howard Hughes Medical Institute, Memorial Sloan Kettering Cancer Center, New York, USA

**Keywords:** YTHDC2, meiosis, pachytene, telomere, chromatin condensation, checkpoint.

## Abstract

Mechanisms driving the prolonged meiotic prophase I are poorly understood. The RNA helicase YTHDC2 is critical for mitosis to meiosis transition, as YTHDC2-deficient mouse germ cells initiate meiosis but arrest with mixed characteristics of mitotic and meiotic cell types. However, YTHDC2 is also highly expressed in normal pachytene cells. Here we identify an essential role for YTHDC2 in meiotic progression. Specifically, we find that YTHDC2 deficiency causes microtubule-dependent telomere clustering and apoptosis at the pachytene stage of prophase I, and thus a failure to advance to the diplotene stage. Depletion of YTHDC2 results in a massively dysregulated transcriptome in pachytene cells, with a tendency toward upregulation of genes normally expressed in mitotic germ cells and downregulation of meiotic transcripts. Dysregulation does not correlate with the m^6^A status of RNAs and YTHDC2-bound mRNAs are enriched in genes upregulated in mutant germ cells, revealing that YTHDC2 primarily targets its substrate mRNAs for degradation. Finally, altered transcripts in YTHDC2-deficient pachytene cells encode microtubule network proteins and inhibition of microtubule polymerization disperses clustered telomeres. Together, our results demonstrate that YTHDC2 regulates the prolonged pachytene stage of prophase I by perpetuating a meiotic transcriptome and preventing changes in the microtubule network that could lead to aberrant telomere clustering.

## INTRODUCTION

In sexually reproducing organisms, meiosis, a cell cycle program unique to germ cells, leads to production of haploid gametes. Germ cells undergo pre-meiotic S-phase DNA replication and initiate meiosis. During meiotic prophase I, homologous chromosomes undergo pairing, synapsis, and recombination (Handel and Schimenti, 2010; Zickler and Kleckner, 2015). At the leptotene stage, synaptonemal complex (SC) proteins such as SYCP2 and SYCP3 begin to assemble along chromosomal axes. Homologous chromosomes pair and begin to synapse, enabled by SC proteins including SYCP1 at the zygotene stage. The pachytene stage is characterized by full synapsis of autosomal homologues and completion of meiotic recombination. Chromosomal desynapsis occurs at the diplotene stage. Meiotic prophase I is generally much longer than mitotic prophase, presumably providing ample time for meiosis-specific chromatin events to complete. In particular, the pachytene stage of meiotic prophase I is the longest, lasting six days in male mice (Russell et al., 1990) and may be required for acquisition of metaphase competence (Guan et al., 2020). However, the temporal maintenance of the prolonged meiotic prophase I remains poorly understood.

*N*^6^-methyladenosine (m^6^A) is the most abundant internal modification in eukaryotic transcripts and a critical regulator of RNA stability, splicing, and translation efficiency (Liu and Pan, 2016; Meyer and Jaffrey, 2014; Yue et al., 2015). In mammals, m^6^A modification is catalyzed by a heterodimer of METTL3 and METTL14 (Bokar et al., 1997; Liu et al., 2014). IME4, a homolog of METTL3, is an inducer of meiosis and mediates N6-adenosine methylation of bulk mRNA exclusively during sporulation in budding yeast, suggesting an ancient role for the modification in the induction of meiosis (Clancy et al., 2002; Schwartz et al., 2013). The functional outcome of m^6^A marks on RNAs is mediated by a family of reader proteins, including YTH domain-containing proteins: YTHDF1, 2, 3 and YTHDC1, 2 (Dominissini et al., 2012; Hsu et al., 2017; Wang et al., 2014; Xu et al., 2014; Xu et al., 2015). Crystal structural analyses show that the YTH domain, an RNA-binding motif, utilizes a conserved aromatic cage to accommodate m^6^A (Xu et al., 2014; Xu et al., 2015; Zhang et al., 2010). Cells deficient for all three YTHDF proteins accumulate the most m^6^A-modified transcripts, showing their combined action on the regulation of mRNA stability (Shi et al., 2017; Zaccara and Jaffrey, 2020).

Unlike other mouse YTH domain m^6^A readers (Kasowitz et al., 2018; Lasman et al., 2020), YTHDC2 is specifically required for meiosis (Bailey et al., 2017; Hsu et al., 2017; Jain et al., 2018; Wojtas et al., 2017). *Ythdc2*-deficient germ cells initiate meiosis but prematurely enter a metaphase state with a mixed mitotic and meiotic identity. In addition to the YTH domain, YTHDC2 contains multiple RNA-binding domains including RNA helicase motifs and ankyrin repeats (Figure S1A). YTHDC2 interacts with the 5′→3′ exoribonuclease XRN1 through the ankyrin repeat (Kretschmer et al., 2018; Wojtas et al., 2017). YTHDC2 also interacts with the small ribosomal subunit (Kretschmer et al., 2018). YTHDC2 is a 3′→5′ RNA helicase (Jain et al., 2018; Wojtas et al., 2017). As YTHDC2 recognizes somatic (mitotic) transcripts such as *Ccna2* for degradation upon meiotic entry, the prevailing model is that YTHDC2 modulates the levels of m^6^A-modified transcripts in meiotic germ cells to generate a transcriptome that facilitates meiotic progression. YTHDC2 partners with MEIOC, a meiosis-specific protein (Abby et al., 2016; Soh et al., 2017). Strikingly, *Ythdc2*-deficient mouse mutants phenocopy the *Meioc*-deficient mutants.

Intriguingly, the YTHDC2 protein is most abundant in mouse pachytene spermatocytes (Bailey et al., 2017; Jain et al., 2018). However, the early meiotic arrest in all four existing *Ythdc2* global mouse mutants precludes investigation of its putative role at the pachytene stage. To circumvent this hurdle, here we employ an inducible inactivation approach coupled with molecular analyses and demonstrate that YTHDC2 is essential for progression through the extended pachytene stage by sustaining a meiotic transcriptome and safeguarding against aberrant microtubule-dependent telomere clustering.

## RESULTS

### Conditional Deletion of *Ythdc2* Leads to Meiotic Blockade at the Pachytene Stage

YTHDC2 is highly expressed in pachytene spermatocytes (Bailey et al., 2017; Jain et al., 2018). To elucidate a potential role at this later stage in meiotic progression, we generated a *Ythdc2* floxed allele by homologous recombination in ES cells (Figure S1B). In this floxed allele, exons 13-16, encoding the ankyrin repeats and part of the HrpA-like RNA helicase domain, are flanked by *loxP* sites. Cre-mediated removal of exons 13-16 results in a deletion of 149 amino acids (aa 578 – 726) and a frameshift in the mutant transcript. We first produced *Ythdc2*^-/-^ (global knockout) mice using the ubiquitous *Actb*-Cre (Lewandoski et al., 1997). *Ythdc2*^-/-^ males exhibited an early meiotic arrest and lacked normal zygotene or later stage spermatocytes, the same phenotype as previously reported (Bailey et al., 2017; Hsu et al., 2017; Jain et al., 2018; Wojtas et al., 2017), showing that this knockout allele is null (Figure S1C). We then conditionally inactivated *Ythdc2* using *Stra8*-Cre, which is primarily expressed in spermatogonia prior to meiotic entry (Baltus et al., 2006; Lin et al., 2017). *Ythdc2*^fl/-^ *Stra8*-Cre conditional knockout males exhibited early meiotic arrest and lacked normal zygotene or later stage spermatocytes (Figure S1D and S1E). Finally, we also conditionally inactivated *Ythdc2* using *Ngn3*-Cre, which begins to express in spermatogonia at postnatal day 7 (Schonhoff et al., 2004). *Ythdc2*^fl/em1^ *Ngn3*-Cre conditional knockout males exhibited a significant reduction in testis size and early meiotic arrest with metaphase-like spermatocytes (Figure S1F and S1G). Therefore, the newly generated *Ythdc2* global knockout and two *Ythdc2* conditional knockout mouse mutants exhibited meiotic defects that were similar to the phenotypes of the previously reported *Ythdc2*^-/-^ mutants.

To circumvent the early meiotic block in both global and conditional knockout *Ythdc2* mutants, we generated *Ythdc2*^fl/-^ *Ddx4*-Cre^ERT2^ mice (referred to as *Ythdc2*^iKO^) for tamoxifen-inducible deletion in germ cells (Figure 1A). *Ddx4*-Cre^ERT2^ is expressed in both mitotic and meiotic germ cells (John et al., 2008). Intraperitoneal tamoxifen injection of adult *Ythdc2*^iKO^ males revealed a progressive decrease in testis weight after treatment (Figure 1B). The abundance of YTHDC2 in *Ythdc2*^iKO^ testes was reduced at 2 days post tamoxifen treatment (2 dpt) and decreased sharply over time (Figure 1C). Histological analysis showed progressively severe spermatogenic defects in *Ythdc2*^iKO^ testes (Figure 1D). Spermatogenesis occurs in waves, and seminiferous tubules are sequentially divided into twelve stages, each of which is defined by a specific cohort of spermatogonia, spermatocytes, and spermatids. Each stage on average lasts about 20 hours in mice (Russell et al., 1990). At 2 dpt, apoptotic pachytene spermatocytes were observed in *Ythdc2*^iKO^ seminiferous tubules (one cell layer away from the basement membrane). Strikingly, these abnormal pachytene cells were present in stages VIII, IX, and X (Figure 1D), but not in early stage tubules (I-VII) (data not shown), and thus were late pachytene cells. In 4 and 6 dpt *Ythdc2*^iKO^ seminiferous tubules, abnormal pachytene cells were still present but pachytene cells were severely depleted in stages VIII-X (Figure S2). In 8 and 10 dpt *Ythdc2*^iKO^ tubules, pachytene spermatocytes were absent (Figure S2). These results revealed a progressive loss of spermatocytes with time after the last tamoxifen injection (Figure S2). Notably, round and/or elongated spermatids from the previous wave of spermatogenesis were still present in *Ythdc2*^iKO^ tubules, further supporting a block in meiotic progression in the current wave.

**Figure 1.**
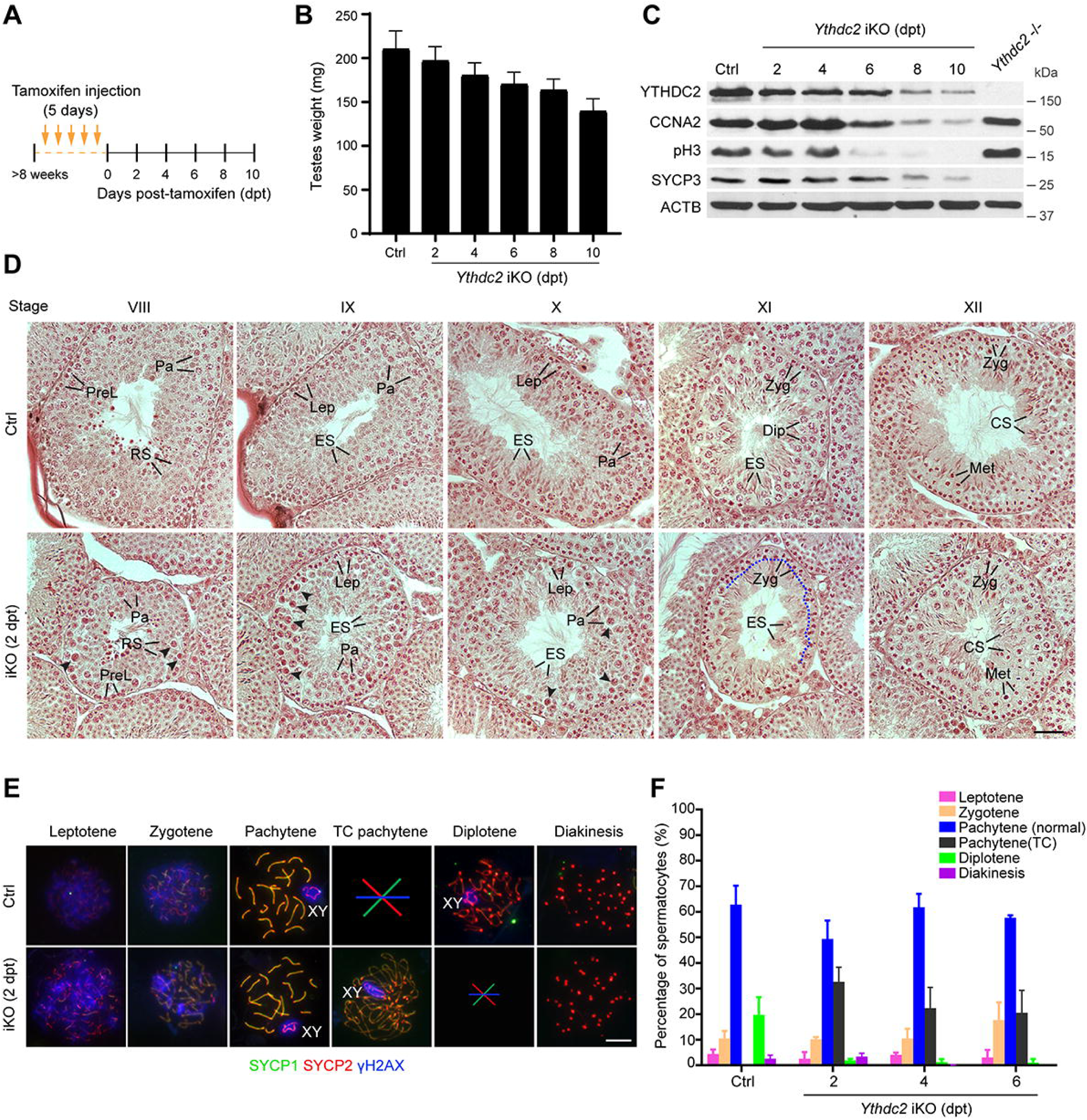
YTHDC2 Is Essential for Progression through the Pachytene Stage. (A) Regimen of tamoxifen treatment in ≥8-week-old *Ythdc2*^fl/-^ *Ddx4*-Cre^ERT2^ male mice. See also Figure S1. (B) Testis weight of control (*Ythdc2*^fl/-^) and *Ythdc2*^iKO^ males (mean ± s.d.; n ≥ 3). (C) Western blot analysis of cell cycle regulators in control (*Ythdc2*^fl/-^), *Ythdc2*^iKO^, and *Ythdc2*^-/-^ testes. (D) Histology of control and *Ythdc2*^iKO^ (2 dpt) testes. Five stages of seminiferous tubules are shown (VIII-XII). Abbreviations: PreL, preleptotene; Lep, leptotene; Zyg, zygotene; Pa, pachytene; Dip, diplotene; Met, metaphase; RS, round spermatid; ES, elongating spermatid; CS, condensing spermatids. Arrowheads indicate apparently apoptotic pachytene spermatocytes. Loss of diplotene spermatocytes at the stage XI *Ythdc2*^iKO^ tubule is demarcated by a dashed blue line. Scale bar, 50 µm. See also Figure S2. (E) Surface nuclear spread analysis of control and *Ythdc2*^iKO^ (2 dpt) spermatocytes. Absence of TC pachytene cells in control is marked with a large symbol. A severe loss of diplotene cells in *Ythdc2*^iKO^ (2 dpt) is indicated by a small symbol. TC, telomere clustering. The percentage of each type of spermatocytes in control and *Ythdc2*^iKO^ testes (mean ± s.d.) is shown in plot (F). >80 cells per mouse and three males per genotype per timepoint were analyzed. Scale bar, 10 µm.

We examined chromosomal synapsis and meiotic recombination in spermatocytes from adult testes by immunofluorescent analysis of SYCP1 (a component of the synaptonemal complex transverse elements), SYCP2 (a component of lateral elements), and γH2AX (Figure 1E). *Ythdc2*^iKO^ pachytene cells displayed normal synapsis as shown by the colocalization of SYCP1 and SYCP2 on all homologues except XY. Meiotic DNA double strand breaks were apparently repaired, since γH2AX was absent on autosomes and restricted to the XY body in *Ythdc2*^iKO^ pachytene cells (Figure 1E).

We next determined the composition of prophase I spermatocytes in adult testes. The control adult testis contained a full spectrum of prophase I spermatocytes from leptotene to diakinesis (Figure 1E and 1F). However, the *Ythdc2*^iKO^ testes contained leptotene, zygotene, and pachytene cells, but were depleted of diplotene cells at 2, 4, and 6 dpt (Figure 1E and 1F). Diplotene spermatocytes are present in stage XI tubules in control, however, histology of *Ythdc2*^iKO^ testes revealed a loss of diplotene spermatocytes at stage XI (Figure 1D and Figure S2). Diakinesis/metaphase I spermatocytes were still present in stage XII tubules from *Ythdc2*^iKO^ testes at 2 dpt (presumably from the previous wave of spermatogenesis), but were absent at 4 dpt and beyond (Figure S2). These results demonstrate that YTHDC2 is required for progression through the pachytene stage of meiotic prophase I.

### Depletion of YTHDC2 Causes Telomere Clustering at the Pachytene Stage

Meiotic chromosome telomeres are attached to the nuclear membrane by the meiotic telomere complex (Shibuya et al., 2015) and connected to the cytoplasmic cytoskeleton network through the LINC complex (Ding et al., 2007; Starr and Han, 2002). In wild type meiotic cells, telomeres are transiently clustered in a phenomenon termed bouquet formation at the zygotene stage, but this clustering no longer occurs during the pachytene stage (Scherthan, 2001; Zickler and Kleckner, 1998). Bouquet formation has been proposed to facilitate homologue search and pairing during meiosis. Unexpectedly, we observed pronounced telomere clustering (termed ‘TC’) as a polarized configuration in pachytene cells from *Ythdc2*^iKO^ testis but not from control testis (Figure 1E). Such abnormal pachytene cells accounted for >20% of prophase I spermatocytes (Figure 1E). In TC pachytene cells, the synaptonemal complexes formed U-shaped loops that had low staining intensity for SC proteins and that appeared to be elongated relative to normal pachytene chromosomes (Figure 1E).

We confirmed telomere clustering in *Ythdc2*^iKO^ pachytene cells with immunofluorescence of CREST (centromere marker) and SUN1 (LINC telomere marker) (Figure 2A) (Crisp et al., 2006; Ding et al., 2007). Mouse chromosomes are acrocentric, i.e., with the centromere positioned at one end. Immunofluorescence of SUN1 in intact *Ythdc2*^iKO^ pachytene cells confirmed telomere clustering (Figure 2B). Confocal microscopy of CREST and SYCP3 showed that centromeres were randomly distributed in *Ythdc2*^fl/-^ pachytene cells (Movie S1) but congregated to one side of the nucleus in *Ythdc2*^iKO^ pachytene cells (Movie S2). To determine when telomeres cluster during the pachytene stage, H1t immunofluorescence was performed. H1t is expressed in mid to late pachytene cells but not early pachytene cells (Figure 2C) (Cobb et al., 1999). *Ythdc2*^iKO^ pachytene cells with telomere clustering were H1t-positive and thus were at the mid-late pachytene stage (Figure 2C). In 2 dpt *Ythdc2*^iKO^ testes, early pachytene cells lacked telomere clustering, nearly half of mid pachytene cells displayed telomere clustering, and most late pachytene cells exhibited telomere clustering (Figure 2D). These results show that *Ythdc2*^iKO^ spermatocytes develop telomere clustering at the mid-late pachytene stage.

**Figure 2.**
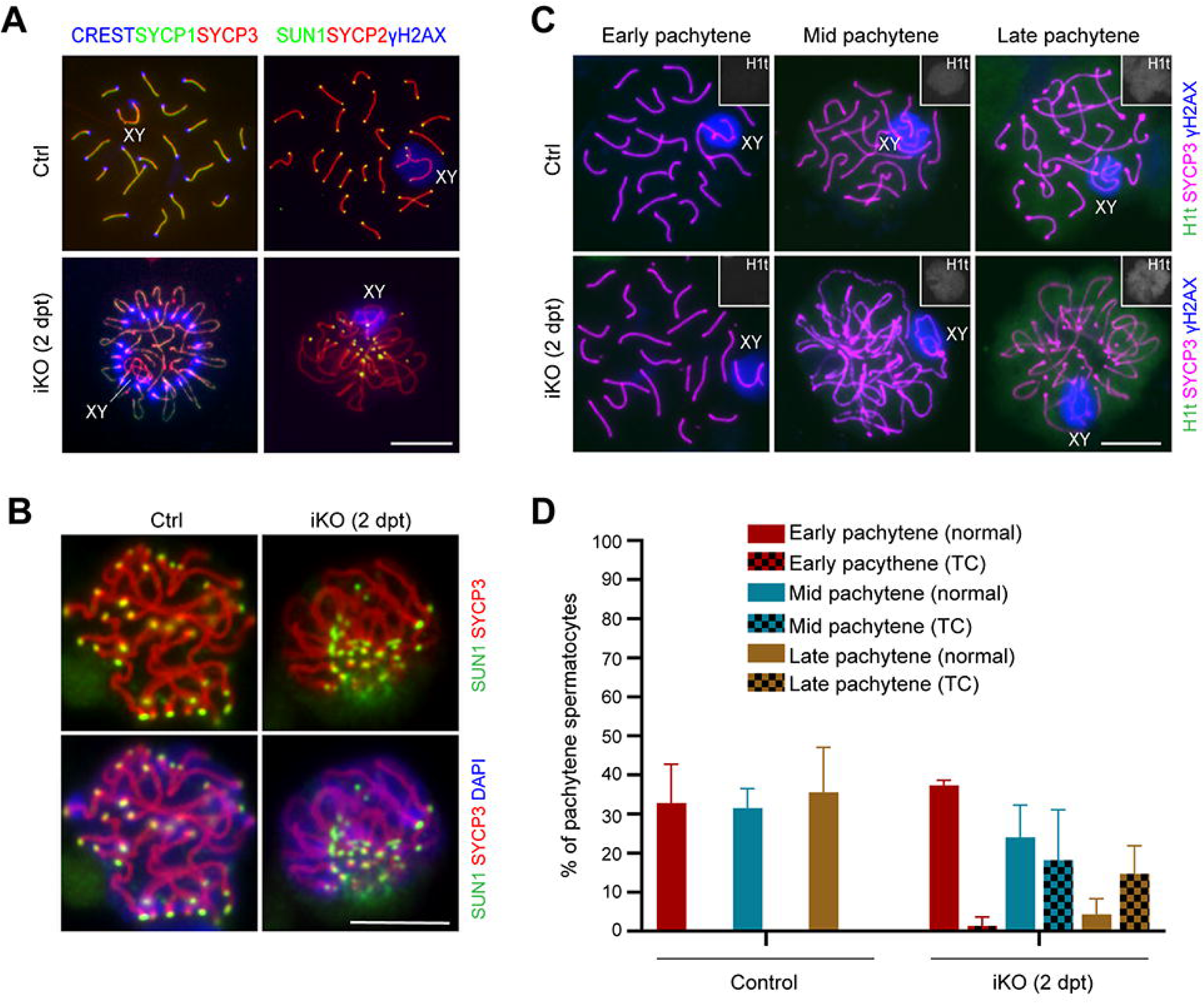
Telomere Clustering Occurs at the Mid-Late Pachytene Stage in *Ythdc2*^iKO^ Spermatocytes. (A) Immunofluorescence of CREST and SUN1 in surface nuclear spreads of pachytene cells. (B) Immunofluorescence of SUN1 in pachytene cells prepared from testicular single cell suspension. See also Movies S1 and S2. (C-D) Substaging of pachytene cells by the H1t expression. H1t is absent in early pachytene cells, appears in mid-pachytene cells, and is abundant in late pachytene cells (C). Inset (C) shows the H1t (black and white) signal in the entire nucleus. The plot (D) shows the percentage (mean ± s.d.) of each type of pachytene spermatocytes. Scale bars, 10 μm.

### Elimination of *Ythdc2*^iKO^ Spermatocytes at the Pachytene Stage

The depletion of diplotene cells in *Ythdc2*^iKO^ testes (Figure 1E) implies that *Ythdc2*^iKO^ pachytene cells die at a specific stage. The pachytene stage of male meiosis is unusually long, spanning six days in mice. To pinpoint the timing of spermatocyte death in *Ythdc2*^iKO^ testes, we performed TUNEL analysis coupled with precise staging of seminiferous tubules (Figure 3). We observed sharply increased apoptosis in *Ythdc2*^iKO^ testes at 2 dpt compared with controls. The occurrence of apoptosis in *Ythdc2*^iKO^ testes was stage-dependent. Tubules at stages VIII, IX, and X displayed the highest percentage (>80%) of tubules with TUNEL-positive cells (Figure 3A and 3B). Stages VIII-X contained late pachytene cells (located one cell layer away from the basement membrane) and pre-leptotene cells (VIII) or leptotene cells (IX and X) (the outermost layer of germ cells) (Figure 3A). Because TUNEL-positive cells in stages VIII-X *Ythdc2*^iKO^ tubules were located one cell layer away from the basement membrane and contained thread-like chromatin (a characteristic of pachytene cells), these apoptotic spermatocytes were late pachytene cells (Figure 3A). Quantification of apoptotic cells in all 12 stages revealed that the level of apoptosis in *Ythdc2*^iKO^ testes at 2 dpt increased significantly at stage VIII and peaked at stages IX and X, whereas the level of apoptosis at stages I – VII and XII remained low in *Ythdc2*^iKO^ testes, similar to control (Figure 3C). In addition, increased apoptosis also occurred in *Ythdc2*^iKO^ testes at 4 and 6 dpt (data not shown). These results demonstrate that mutant spermatocytes are specifically eliminated by apoptosis at the late pachytene stage, resulting in the absence of diplotene cells in *Ythdc2*^iKO^ testis.

**Figure 3.**
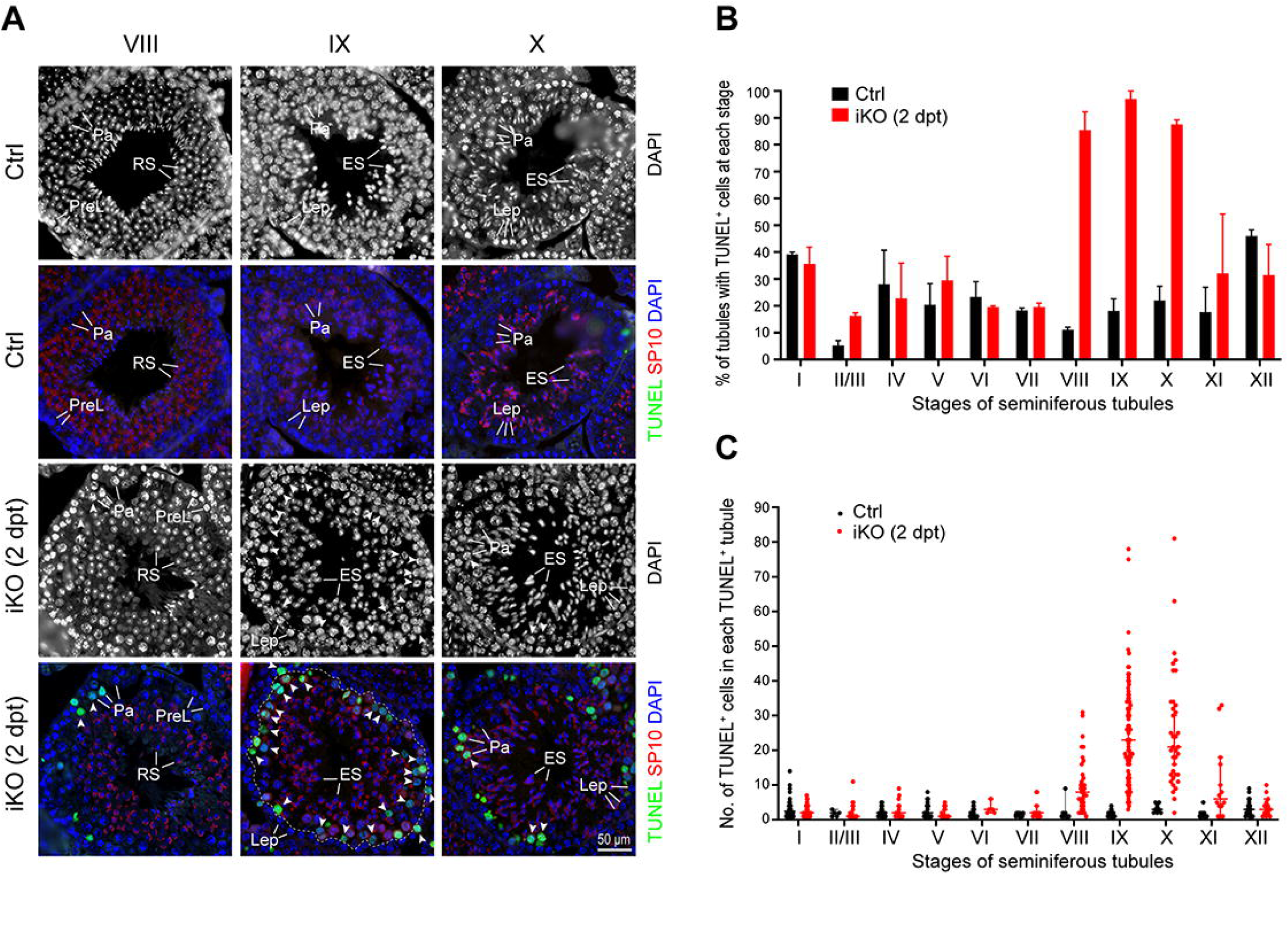
*Ythdc2*^iKO^ Spermatocytes Undergo Apoptosis at the Late Pachytene Stage. (A) TUNEL analysis of paraffin-embedded tissue sections from control and *Ythdc2*^iKO^ (2 dpt) testes. The acrosome morphology shown by SP10 immunofluorescence staining was used for seminiferous tubule staging. Nuclear DNA was stained with DAPI. Arrowheads indicate TUNEL-positive pachytene cells. The dashed line in stage IX *Ythdc2*^iKO^ demarcates pachytene cells (the inner layer) from leptotene cells (the outer layer). Abbreviations: PreL, preleptotene; Lep, leptotene; Pa, pachytene; RS, round spermatid; ES, elongating spermatid. Scale bar, 50 μm. (B) Percentage of TUNEL-positive tubules from control and *Ythdc2*^iKO^ testes at 2 dpt. The mean ± s.d. values were plotted. (C) Quantification of TUNEL-positive cells in TUNEL-positive tubules (mean ± s.d.) from control and *Ythdc2*^iKO^ at 2 dpt. (B, C) Two males per genotype (control and *Ythdc2*^iKO^) were analyzed. At least 200 tubules were counted for each mouse.

### *Ythdc2*^iKO^ Leptotene Cells Prematurely Condense Chromatin

Male germ cells (preleptotene cells) in *Ythdc2*-deficient mice initiate meiosis and enter prematurely into a mitotic metaphase state with condensed chromatin (Bailey et al., 2017; Hsu et al., 2017; Jain et al., 2018; Wojtas et al., 2017). To investigate whether this is the case for *Ythdc2*^iKO^ male germ cells, we performed immunofluorescence analysis of pH3 (phosphorylated histone H3 at serine 10), which is a metaphase marker (Sun and Handel, 2008). In control testis, metaphase I/II spermatocytes (pH3-positive) were only present at stage XII tubules (Figure S3A). Likewise, in 2 dpt *Ythdc2*^iKO^ testis, most pH3-positive spermatocytes were observed at stage XII, and the preleptotene (stage VIII) and leptotene (stages IX and X) were pH3-negative (Figure S3A and S3B). Strikingly, in 4 dpt *Ythdc2*^iKO^ testis, the outermost layer of germ cells at stages VIII, IX, and X were pH3-positive and exhibited condensed chromatin, indicating a metaphase-like state (Figure S3A). Quantification of pH3-positive cells showed the highest number of pH3-positive cells at stage IX tubules from 4 dpt *Ythdc2*^iKO^ testis (Figure S3C). In addition, 6 dpt *Ythdc2*^iKO^ testis contained pH3-positive metaphase-like germ cells at stages VIII, IX, and X (Figure S3A). Consistently, histological analyses showed that leptotene cells and zygotene cells were present in *Ythdc2*^iKO^ tubules at 2 dpt and 4 dpt, but severely depleted at 6 dpt (Figure S2). Furthermore, zygotene cells were absent in stage XI/XII *Ythdc2*^iKO^ tubules at 8 and 10 dpt (Figure S2). These results demonstrate that *Ythdc2*^iKO^ male germ cells enter meiosis but fail to progress through the leptotene stage and prematurely condense chromatin in a metaphase-like state.

### *Ythdc2*^iKO^ Pachytene Cells with Telomere Clustering Are Metaphase-Competent

Okadaic acid, an inhibitor of protein phosphatases PP1 and PP2A, induces progression of pachytene cells into metaphase I stage *in vitro* (Wiltshire et al., 1995). *Ythdc2*^iKO^ pachytene cells failed to progress to the diplotene stage in the testis. To address whether *Ythdc2*^iKO^ pachytene cells with telomere clustering are competent for metaphase I entry, spermatocytes were treated with okadaic acid in culture (Figure 4A). Among both control (*Ythdc2*^fl/-^) and *Ythdc2*^iKO^ spermatocytes, the percentage of diakinesis/metaphase I spermatocytes increased to over 80% after OA treatment (Figure 4B). Accordingly, the percentage of *Ythdc2*^iKO^ pachytene cells with telomere clustering was sharply reduced. These results demonstrate that *Ythdc2*^iKO^ pachytene cells are competent for entry into metaphase I.

**Figure 4.**
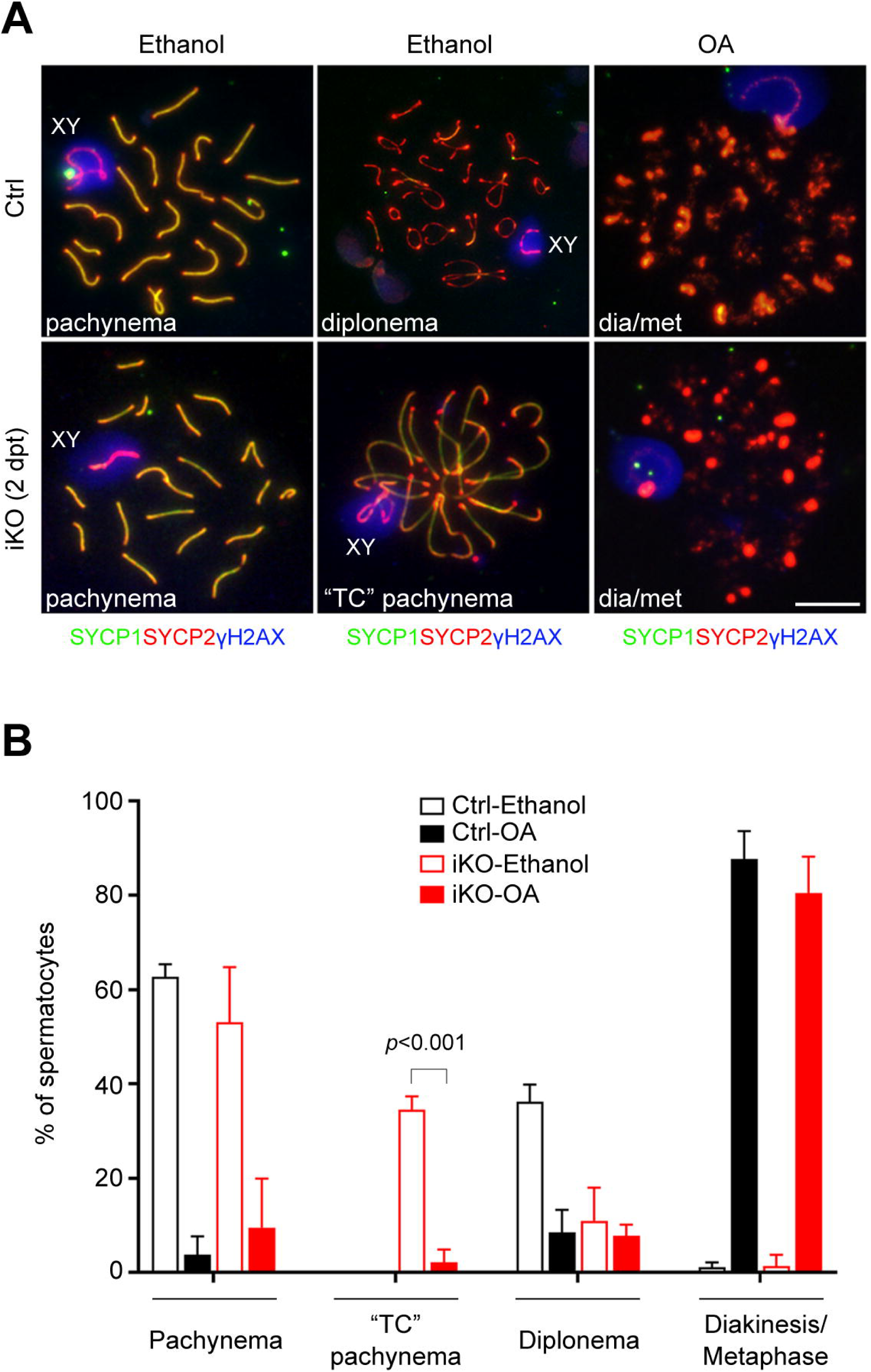
Meiotic progression of *Ythdc2*^iKO^ pachytene cells with telomere clustering by okadaic acid (OA) treatment. (A) Images of pachytene, diplotene, and diakinesis/metaphase I (dia/met) spermatocytes before and after OA treatment. Scale bar, 10 μm. (B) Percentage of spermatocytes before and after OA treatment (mean ± s.d.). Only pachytene, diplotene, and diakinesis/metaphase I spermatocytes are included in this analysis. At least 250 cells were counted per genotype per treatment group. Two males per genotype per experiment were used. The experiments were performed three times (n=3).

### Massively Altered Transcriptome in *Ythdc2*^iKO^ Pachytene Cells

To elucidate the molecular defects in *Ythdc2*^iKO^ pachytene spermatocytes, we performed RNA-seq analysis. Because whole testes contained both somatic and germ cells and differed in cellular composition between control and mutant, pachytene cells were purified by the Sta-put method and used for RNA-seq analysis (Figure S4A). A large number of differentially expressed genes were identified between *Ythdc2*^fl/-^ (control) and *Ythdc2*^iKO^ pachytene cells at 2 dpt (Figure 5A). With the cutoff of log_2_(Fold Change) ≥ 0.5 or ≤ -0.5, RNA-seq analysis revealed more up-regulated genes (3147 genes) than down-regulated genes (1812 genes) in *Ythdc2*^iKO^ pachytene cells (Figure 5B and Table S1). The expression changes of a set of 16 differentially expressed genes (8 upregulated and 8 downregulated) were validated and confirmed by qPCR (Figure S4B and S4C). Western blot analysis of purified pachytene cells showed that the abundance of YTHDC2 was reduced in *Ythdc2*^iKO^ pachytene cells, as expected (Figure S4D). Consistent with the downregulation of its transcript, SYCP2 protein abundance was reduced in *Ythdc2*^iKO^ pachytene cells (Figure S4D). Even though the *Meioc* transcript was significantly increased in the *Ythdc2*^iKO^ cells, the abundance of MEIOC protein remained constant (Figure S4D), potentially because the reduced abundance of its binding partner YTHDC2 in *Ythdc2*^iKO^ cells led to its instability (Figure S5A).

**Figure 5.**
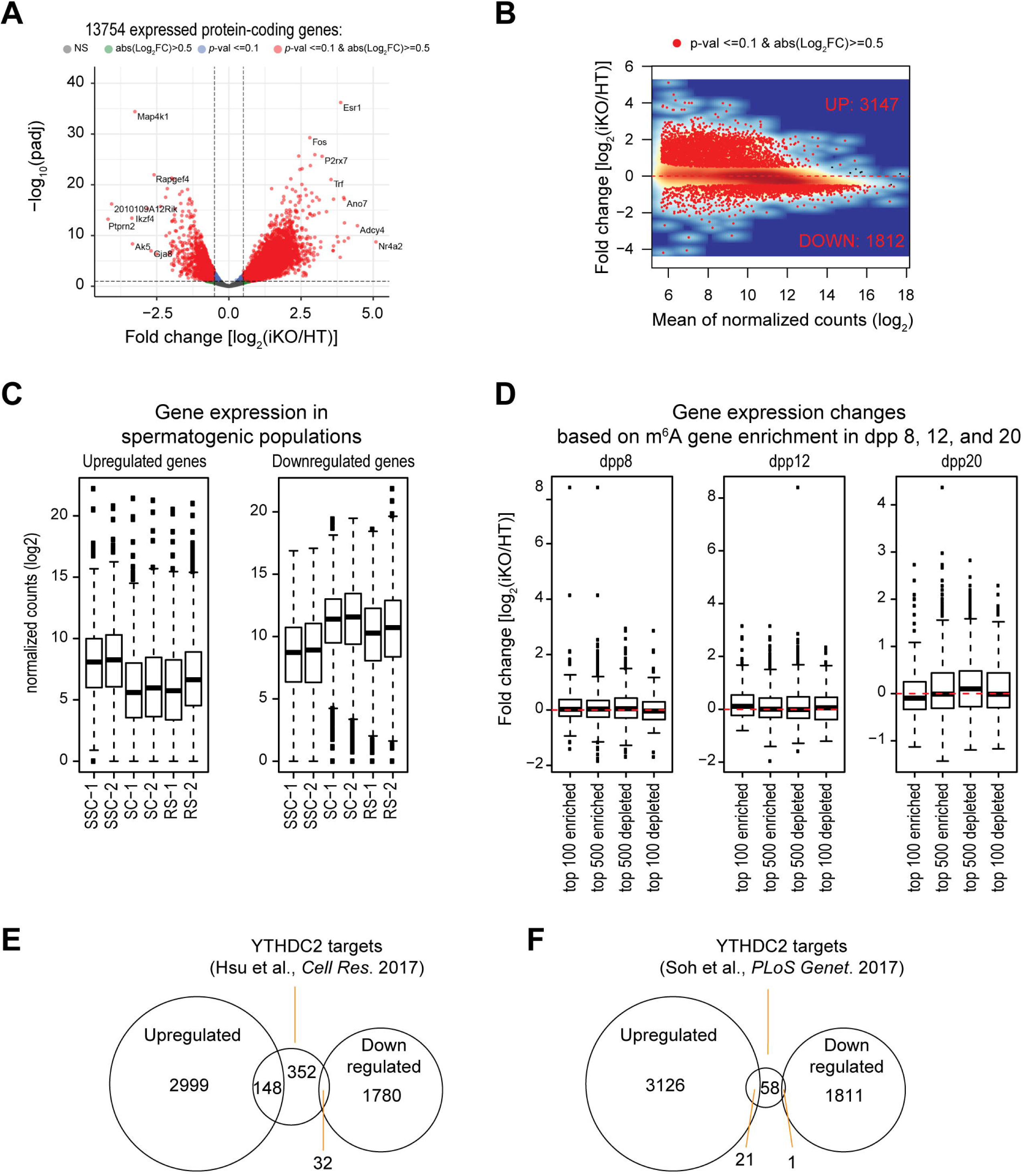
Dramatic alteration of transcriptome in *Ythdc2*^iKO^ pachytene spermatocytes. (A) Volcano plot of expression changes in *Ythdc2*^iKO^ vs control (*Ythdc2*^fl/-^; HT) pachytene spermatocytes. Only coding genes (13,754) with an average expression of ≥50 across the samples are plotted. (B) MA plot of expression changes. The differentially expressed (up and down regulated) genes are shown as red dots. See also Figures S4 and S5. (C) Expression of differentially expressed genes in various wild-type spermatogenic populations. SSC, spermatogonial stem cells; SC, spermatocytes; RS, round spermatids. (D) Expression comparison of genes in *Ythdc2*^iKO^ vs control (*Ythdc2*^fl/-^; HT) pachytene cells that are enriched or depleted of m^6^A in juvenile testes (Wojtas et al., 2017). (E) Overlap of YTHDC2 RIP-seq targets (Hsu et al., 2017) with the differentially expressed genes in *Ythdc2*^iKO^ pachytene cells. (F) Overlap of YTHDC2 RIP-seq targets (Soh et al., 2017) with the differentially expressed genes in *Ythdc2*^iKO^ pachytene cells.

The top enriched GO terms for upregulated genes are actin binding, focal adhesion, regulation of signal transduction, and regulation of apoptosis (Figure S5B). The top enriched GO terms for downregulated genes are RNA-binding, microtubule organizing center, and cilium assembly and organization (Figure S5C). We next examined the expression profile of differentially expressed genes among specific spermatogenic populations: spermatogonia (mitotic), spermatocytes (meiotic), and round spermatids (post-meiotic), using the germ cell RNA-seq data (Davis et al., 2017). The upregulated genes in *Ythdc2*^iKO^ pachytene cells exhibit higher expression levels in spermatogonia than spermatocytes, suggesting that YTHDC2 promotes degradation of mitotic transcripts in spermatocytes (Figure 5C). In contrast, the downregulated genes normally show higher expression levels in spermatocytes (Figure 5C).

Notably, the overall transcript levels of only X chromosome-linked genes and mitochondria-encoded genes were upregulated in *Ythdc2*^iKO^ pachytene cells (Figure S5D). Although all autosomes had both upregulated and downregulated genes, all differentially expressed X-linked genes (274 genes) were upregulated (Figure S5E).

### YTHDC2 Destabilizes Its Target Transcripts

YTHDC2-bound transcripts are enriched for the m^6^A modification (Bailey et al., 2017; Hsu et al., 2017; Wojtas et al., 2017). We analyzed the *Ythdc2*^iKO^ expression changes among genes known to be enriched or depleted of m^6^A (Wojtas et al., 2017), but found no correlation between expression changes in *Ythdc2*^iKO^ pachytene cells and the level of m^6^A in mRNAs (Figure 5D). This could indicate that the m^6^A-binding activity of YTHDC2 is dispensable or that these expression changes are mostly indirect effects.

YTHDC2 contains multiple RNA-binding domains: YTH, R3H, and RNA helicase domain (Figure S1A) (Jain et al., 2018; Kretschmer et al., 2018; Wojtas et al., 2017). YTHDC2 lacking the YTH domain still binds to cellular RNAs (Kretschmer et al., 2018). The target mRNAs of YTHDC2 in testes were previously identified by RNA immunoprecipitation sequencing (RIP-seq) and 521 YTHDC2 target mRNAs were reported (Hsu et al., 2017). Out of these 521 genes, 148 genes (28%) were upregulated, but only 32 genes (6%) were downregulated in *Ythdc2*^iKO^ pachytene cells (Figure 5E and Table S2). Out of 80 YTHDC2 target transcripts reported in another study (Soh et al., 2017), 21 genes (26%) were upregulated but only one was downregulated in *Ythdc2*^iKO^ pachytene cells (Figure 5F and Table S2). Both *Pbrm1* and *Rad21* are YTHDC2 target transcripts identified by RIP-seq (Table S2) and are upregulated in *Ythdc2*^iKO^ pachytene cells (Table S1). PBRM1 is a component of the PBAF chromatin-remodeling complex (Brownlee et al., 2014) and RAD21 is a mitotic/meiotic cohesin (Xu et al., 2004). The protein abundance of PBRM1 and RAD21 was increased in *Ythdc2*^iKO^ pachytene cells (Figure S4D). In conclusion, the preferential upregulation of YTHDC2 target transcripts in *Ythdc2*^iKO^ pachytene cells further supports that YTHDC2 primarily targets transcripts for degradation.

### Microtubule-Dependent Telomere Clustering in *Ythdc2*^iKO^ Pachytene Cells

The top GO terms of downregulated genes in *Ythdc2*^iKO^ pachytene cells are related to microtubules: microtubule organizing center, centriole, and cilium (Figure S5C). Even though microtubule-related categories are not among the top 10 GO terms for upregulated genes (Figure S5B), 39 upregulated genes encode tubulins, microtubule-associated proteins, dyneins, and kinesins (Table S1). In mammalian cells, telomere-led meiotic chromosome movement is primarily driven by cytoplasmic microtubule-based forces (Lee et al., 2015). To address whether the microtubule cytoskeleton is responsible for abnormal telomere clustering, we examined the microtubule network in intact pachytene cells. In *Ythdc2*^fl/-^ pachytene cells, cytoplasmic microtubules were randomly distributed and telomeres (indicated by SUN1) were distributed throughout the nuclear periphery (Figure 6A). Strikingly, in 83% of *Ythdc2*^iKO^ pachytene cells with clustered telomeres (83±8%; 82 TC pachytene cells from three males were analyzed), microtubules converged to the region where telomeres were clustered (Figure 6A). In addition, the side where both microtubules and telomeres congregated was expanded cytoplasm (Figure S6). These results suggest that telomere clustering could be attributed to the polarized distribution of cytoplasmic microtubules in *Ythdc2*^iKO^ pachytene cells.

**Figure 6.**
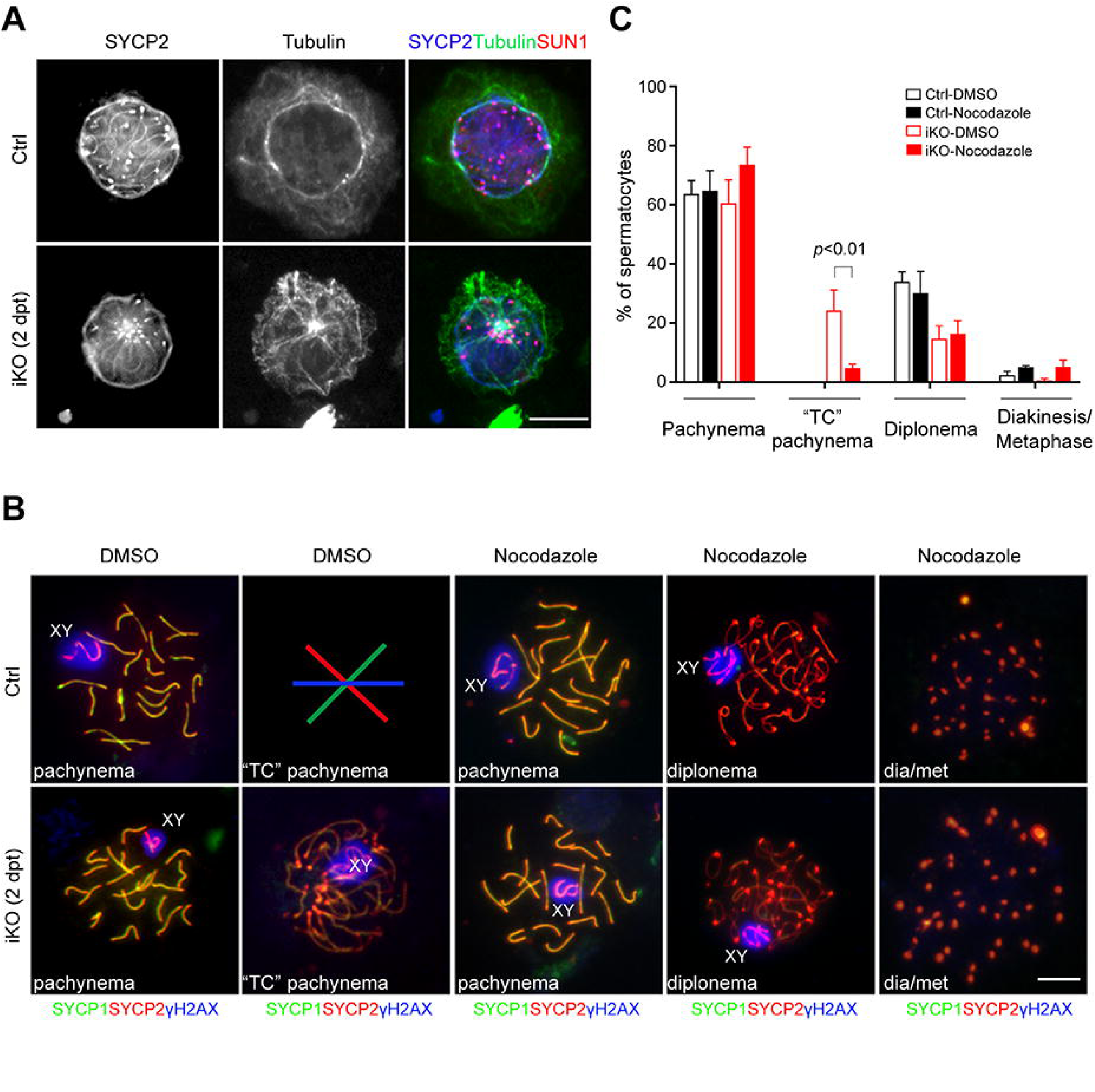
Inhibition of microtubule polymerization rescues telomere clustering in *Ythdc2*^iKO^ pachytene cells. (A) Immunofluorescence of α-tubulin, SUN1, SYCP2 in intact control (*Ythdc2*^fl/-^) and *Ythdc2*^iKO^ (2 dpt) pachytene spermatocytes. Scale bar, 10 μm. See also Figure S6. (B) Nuclear spread analysis of pachytene, diplotene, and diakinesis/metaphase I (dia/met) spermatocytes after treatment with DMSO or nocodazole. TC, telomere clustering. Scale bar, 10 μm. (C) Percentage of spermatocytes (mean ± s.d.) in control (*Ythdc2*^fl/-^) and *Ythdc2*^iKO^ (2 dpt) after treatment with DMSO or nocodazole. Only pachytene, diplotene, and diakinesis/metaphase I spermatocytes are included. At least 500 cells were counted per genotype per treatment group. Two males per genotype per experiment were used. The experiments were performed three times (n=3). See also Figure S7.

To further test the role of microtubules in telomere clustering, *Ythdc2*^iKO^ germ cells were treated with nocodazole, an inhibitor of microtubule polymerization (Figure 6B). The percentage of *Ythdc2*^iKO^ pachytene cells with telomere clustering was significantly reduced from 24.2% (24.2 ± 6.9%, n= 3, *p* < 0.01) in DMSO treatment to 4.8% (4.8 ± 1.3%) in nocodazole treatment (Figure 6C). Treatment of *Ythdc2*^iKO^ cells with colcemid, another microtubule inhibitor, also reduced the percentage of pachytene cells with telomere clustering (data not shown). The top GO terms of the upregulated genes in *Ythdc2*^iKO^ pachytene cells are related to actin: actin-binding and focal adhesion (Figure S5B). However, treatment with actin inhibitors (latrunculin A and cytochalasin D) did not significantly affect the percentage of *Ythdc2*^iKO^ pachytene cells with telomere clustering (Figure S7). Collectively, these results demonstrate that inhibition of microtubule but not actin polymerization rescues abnormal telomere clustering in *Ythdc2*^iKO^ pachytene cells.

## DISCUSSION

All previous studies of *Ythdc2* global knockout mice report a requirement for YTHDC2 around the point of meiotic entry (Figure 7A) (Bailey et al., 2017; Hsu et al., 2017; Jain et al., 2018; Wojtas et al., 2017). In this study, we employed an inducible inactivation approach and uncovered an additional novel function for YTHDC2 in meiosis – progression through the extended pachytene stage during meiotic prophase I. This new finding is consistent with the fact that YTHDC2 is most abundant in pachytene cells. Therefore, YTHDC2 plays essential roles at two distinct stages of meiotic prophase I: at the time of meiotic entry and during late pachytene (Figure 7A). At the early point, YTHDC2-deficient spermatocytes fail to downregulate mitotic genes such as *Ccna2* but undergo chromatin condensation with mixed mitotic/meiotic characteristics (Bailey et al., 2017; Hsu et al., 2017; Jain et al., 2018; Wojtas et al., 2017). Our *Ythdc2*^iKO^ study confirms the requirement for YTHDC2 at this early stage and, furthermore, pinpoints the time of chromatin condensation (pH3-positivity) to seminiferous tubule stages VIII-IX, which is the time when spermatocytes would normally be in pre-leptotene and leptotene stages. Thus, YTHDC2 prevents leptotene cells from reverting to a mitotic cell cycle program. At the pachytene stage, depletion of YTHDC2 causes a massively dysregulated transcriptome. *Ythdc2*^iKO^ pachytene cells are arrested at the late pachytene stage and fail to progress to the diplotene stage. Unlike *Ythdc2*-deficient leptotene cells, *Ythdc2*^iKO^ pachytene cells are pH3-negative and do not undergo chromatin condensation, suggesting that pachytene cells have irreversibly committed to the meiotic program.

**Figure 7.**
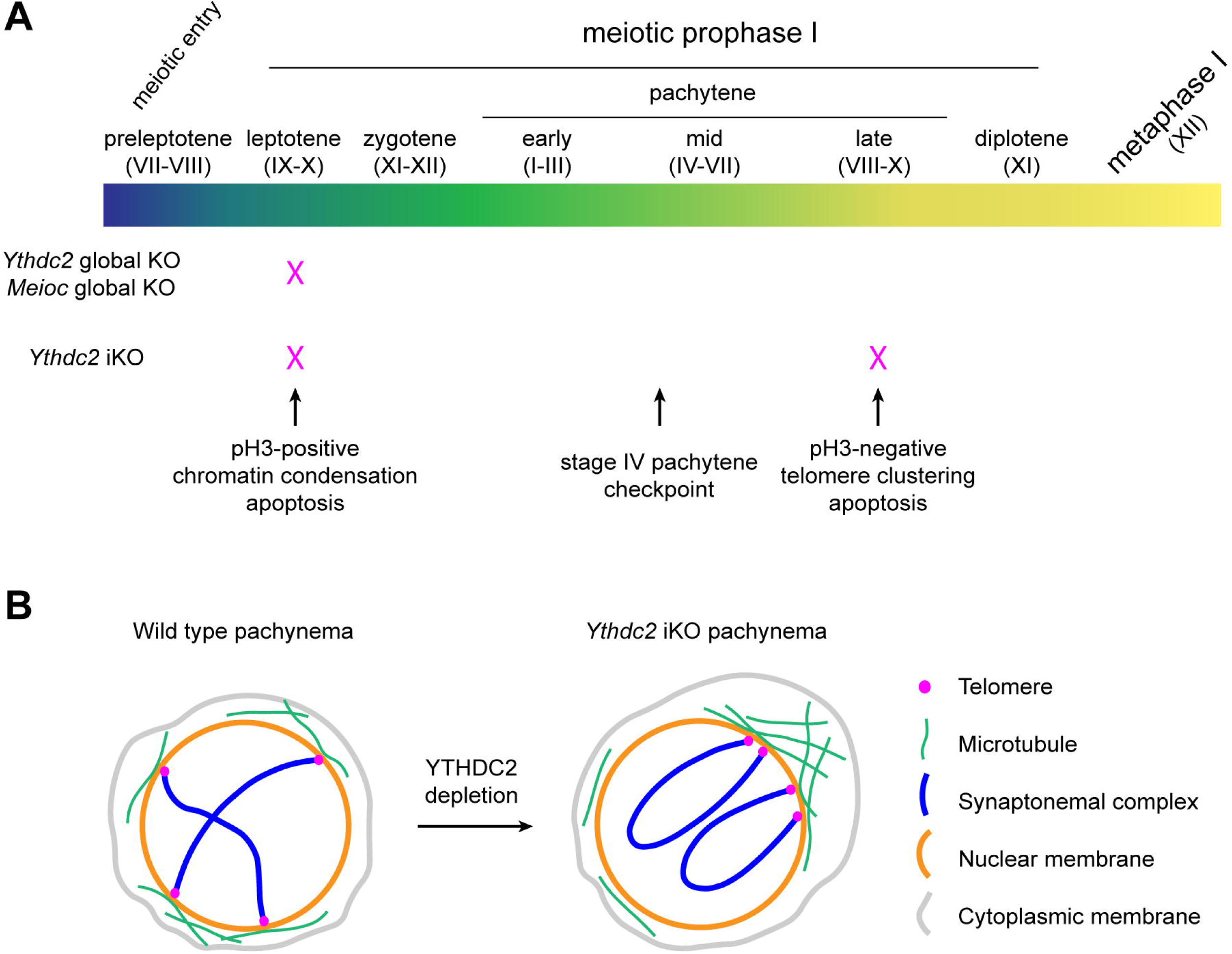
Stage-Specific Regulation of Meiotic Prophase I Progression by YTHDC2. (A) Requirement of YTHDC2 at the leptotene and pachytene stages. The red cross designates the stage of meiotic arrest in the *Ythdc2* or *Meioic* mutants. *Ythdc2* or *Meioc* global knockout mutants display the same early meiotic arrest, which is also present in the *Ythdc2*^iKO^ (*Ythdc2*^fl/-^ *Ddx4*-Cre^ERT2^) mutant. In contrast, *Ythdc2*^iKO^ pachytene cells undergo apoptosis at the late pachytene stage. The seminiferous tubule stage is shown in Roman numerals. See also Figure S3. (B) Illustration of telomere distribution in wild type and *Ythdc2*^iKO^ pachytene spermatocytes. Two pairs of homologous chromosomes are depicted. The SUN/KASH proteins that connect telomeres with microtubules are not shown. In contrast with the random distribution of telomeres in wild type, telomeres in the *Ythdc2*^iKO^ pachytene cell are clustered to one pole, where microtubules converge and the cytoplasm is expanded.

Defects in chromosomal synapsis and meiotic recombination lead to mid-pachytene arrest at stage IV in males, presumably due to a pachytene checkpoint (de Rooij and de Boer, 2003). Although the pachytene checkpoint in yeast is well known (Roeder and Bailis, 2000), components of the pachytene checkpoint in mice have not been identified. XY chromosomes undergo MSCI (meiotic sex chromatin inactivation) during meiosis (Turner, 2015) and genetic studies demonstrate that an MSCI checkpoint operates at stage IV (Abe et al., 2020; Royo et al., 2010). In early pachytene cells, DNA damage repair factors are redistributed from autosomes to XY to achieve MSCI (Abe et al., 2020). Ectopic expression of two Y-linked genes *Zfy1* and *Zfy2* is sufficient to cause death of pachytene spermatocytes at stage IV, suggesting that escape of *Zfy1/2* from MSCI contributes mechanistically to the MSCI checkpoint (Royo et al., 2010). In this study, we report for the first time the death of *Ythdc2*^iKO^ meiotic cells at the late pachytene stage (VIII-X). The late pachytene apoptosis of *Ythdc2*^iKO^ meiotic cells could be due to the dramatically altered transcriptome in *Ythdc2*^iKO^ pachytene cells or to the existence of a late pachytene checkpoint. The transcript levels of *Zfy1* and *Zfy2* remain unchanged in *Ythdc2*^iKO^ pachytene cells and thus are not responsible for apoptosis. However, 274 X-linked genes were upregulated but none were downregulated in *Ythdc2*^iKO^ pachytene cells and *Zfx*, the X-linked paralogue of *Zfy1/2* (Luoh et al., 1997), was upregulated in *Ythdc2*^iKO^ pachytene cells (Table S1). Thus, it is possible that overexpression of X-linked genes such as *Zfx* might be lethal to late pachytene cells.

Telomeres are connected to the cytoplasmic cytoskeleton network through the SUN/KASH proteins (Crisp et al., 2006; Starr and Han, 2002). Telomere clustering (the so-called bouquet formation) occurs only in zygotene cells in wild type (Scherthan, 2001). Telomere-led rapid chromosome movement driven by cytoplasmic forces also occurs in pachytene cells and its function is likely to resolve chromosome entanglements (Lee et al., 2015). Unexpectedly, *Ythdc2*^iKO^ cells exhibit abnormal telomere clustering at the late pachytene stage. This defect is caused by dysregulated cytoplasmic microtubule network but not actin cytoskeleton (Figure 7B). This conclusion is supported by three lines of evidence. First, a large number of differentially expressed (both upregulated and downregulated) genes in *Ythdc2*^iKO^ pachytene cells encode tubulins, microtubule-associated proteins, dyneins, and kinesins (Table S1). Second, cytoplasmic microtubules and telomeres converge to the same pole in *Ythdc2*^iKO^ pachytene cells where the cytoplasm is expanded. This observation supports a direct connection between polarized microtubule networks and telomere clustering. Finally, telomere clustering in *Ythdc2*^iKO^ cells can be resolved by treatment with inhibitors of microtubules but not actin filaments.

YTHDC2 harbors multiple RNA-binding domains, including the m^6^A-recognizing YTH domain and RNA helicase domains. YTHDC2 is an obligate binding partner of MEIOC and *Ythdc2* global knockout mutants phenocopies *Meioc* global knockout mutants (Abby et al., 2016; Bailey et al., 2017; Hsu et al., 2017; Jain et al., 2018; Soh et al., 2017; Wojtas et al., 2017). Like YTHDC2, MEIOC is abundantly expressed in pachytene cells (Abby et al., 2016; Soh et al., 2017). Therefore, we postulate that inducible inactivation of MEIOC is likely to cause the same defects as *Ythdc2*^iKO^: abnormal telomere clustering in pachytene cells and meiotic arrest at the late pachytene stage. MEIOIC lacks known RNA-binding domains and thus the exact mechanism of MEIOC function is unknown. YTHDC2 interacts with XRN1, a 5′→3′ exoribonuclease (Kretschmer et al., 2018; Wojtas et al., 2017). The YTHDC2/MEIOC complex targets its mitotic substrate transcripts for degradation (Soh et al., 2017; Wojtas et al., 2017). In support of the RNA degradation model, the reported YTHDC2 target transcripts are preferentially upregulated in *Ythdc2*^iKO^ pachytene cells. In addition, we find that differential gene expression in *Ythdc2*^iKO^ pachytene cells does not correlate with the m^6^A RNA modification, suggesting that YTHDC2 might modulate the transcriptome independently of the m^6^A modification. In conclusion, our study demonstrates that YTHDC2 and presumably MEIOC as well continue to be required for actively maintaining a meiotic transcriptome at the prolonged pachytene stage of meiotic prophase I long after meiotic entry.

## ACKNOWLEDGMENTS

We thank David Page for MEIOC antibody and Leslie King for help with manuscript preparation. We thank Jessica Chotiner, Yongjuan Guan, and Fang Yang for critical reading of the manuscript. This work was supported by National Institutes of Health/National Institute of General Medical Sciences grant R35GM118052 (PJW), China Scholarship Council fellowship (201906275011) (RL), the Swiss National Science Foundation via the NCCR RNA & Disease network (51NF40_182880) (RSP), National Key Research & Development Program of China (2018YFC1003400) (ML), the Howard Hughes Medical Institute (SK), and a fellowship from the Human Frontier Science Program (DJ).

## AUTHOR CONTRIBUTIONS

PJW, RL, and SDK conceptualized the study and designed the experiments. RL performed most of the experiments. ML identified YTHDC2. SDK generated the *Ythdc2 floxed* mice. NAL performed blastocyst injection of targeted ES cells. SDK and JS generated the *Ythdc2^fl/-^ Ddx4-Cre^ERT2^* mice. DH and RSP analyzed the RNA-seq data. GR contributed to microscopy and performed 3D deconvolution. DJ and SK generated the *Ythdc2*^fl/em1^ *Ngn3*-Cre data. PJW and RL wrote the manuscript.

## DATA AND MATERIALS AVAILABILITY

All data needed to evaluate the conclusions in the paper are present in the paper and/or the Supplementary Materials. The original RNA-seq data of the current research has been deposited to GEO with accession number GSE166568.

## STAR * METHODS

### Key Resources Table

| REAGENT or RESOURCE | SOURCE | IDENTIFIER |
| --- | --- | --- |
| **Antibodies** |  |  |
| Mouse anti-ACTB | Sigma | Cat# A5441, RRID: RRID:AB_476744 |
| Rabbit anti-CCNA2 | Abcam | Cat# ab181591, RRID:AB_2890136 |
| Human anti-centromere (CREST) | Antibodies Incorporated | Cat# 15-234, RRID:AB_2687472 |
| Guinea pig anti-H1t | Custom-made, a gift from Mary Ann Handel (Cobb et al., 1999) | N/A |
| Rabbit anti-MEIOC | Custom-made, a gift from David Page (Soh et al., 2017) | N/A |
| Rabbit anti-pH3 | Cell Signaling Technology | Cat# 9701S, RRID:AB_331535 |
| Guinea pig anti-SP10 (ACRV1) | Custom-made, a gift from Prabhakara Reddi (Reddi et al., 1995) | N/A |
| Rabbit anti-SUN1 | Abcam | Cat# ab103021, RRID:AB_2890137 |
| Rabbit anti-SYCP1 | Abcam | Cat# ab15090, RRID:AB_301636 |
| Guinea pig anti-SYCP2 | Custom-made, (Yang et al., 2006) | N/A |
| Mouse anti-SYCP3 | Abcam | Cat# ab97672, RRID:AB_10678841 |
| Rabbit anti-SYCP3 | ProteinTech Group | Cat# 23024-1-AP, RRID:AB_11232426 |
| Rabbit anti-YTHDC2 | Novus Biologicals | Cat# NBP1-85089, RRID:AB_11034900 |
| Rabbit anti-YTHDC2 | Custom made, this study | UP2563 |
| Mouse anti-α-Tubulin-FITC antibody | Sigma | Cat# F2168, RRID:AB_476967 |
| Mouse anti-γH2AX | Millipore | Cat# 16-202A, RRID:AB_568825 |
| **Chemicals** |  |  |
| Tamoxifen | Sigma | Cat# T5648-1G |
| Okadaic acid | Cell Signaling Technology | Cat# 5934S |
| Nocodazole | Santa Cruz Biotechnology | Cat# sc-3518C |
| Colcemid | Sigma | Cat# 10295892001 |
| Latrunculin A | Sigma | Cat# 428021-100UG |
| Cytochalasin D | Sigma | Cat# C8273-1MG |
| **Deposited Data** |  |  |
| *Ythdc2* pachytene RNA-seq data | This study | GEO: GSE166568 |
| m<sup>6</sup>A_seq data from mouse testes | (Wojtas et al., 2017) | GEO: GSE102346 |
| RNA_seq profiling of mouse germ cells | (Davis et al., 2017) | ENA: PRJEB15333 |
| **Experimental Models: Organisms/Strains** |  |  |
| Mouse: *Actb*-Cre | The Jackson Laboratory (Lewandoski et al., 1997) | Stock No: 003376<br>RRID:IMSR_JAX:003376 |
| Mouse: *Stra8*-Cre | (Lin et al., 2017) | N/A |
| Mouse: *Neurog3*-Cre | The Jackson Laboratory (Schonhoff et al., 2004) | Stock No: 006333<br>RRID:IMSR_JAX:006333 |
| Mouse: *Ddx4*-Cre<sup>ERT2</sup> | The Jackson Laboratory (John et al., 2008) | Stock No: 024760<br>RRID:IMSR_JAX:024760 |
| Mouse: *Ythdc2* fl/fl | This study | N/A |
| **Software and Algorithms** |  |  |
| Adobe Creative Cloud | Adobe | https://www.adobe.com/uk/creativecloud.html |
| Fiji-ImageJ | (Schneider et al., 2012) | https://imagej.nih.gov/ij/ |
| GraphPad Prism 8.0 | GraphPad | https://www.graphpad.com |
| SPSS Statistics 26 | IBM | https://www.ibm.com/products/spss-statistics |

## CONTACT FOR REAGENT AND RESOURCE SHARING

Further information and requests for resources and reagents should be directed to and will be fulfilled by the Lead Contact, P. Jeremy Wang.

### Targeted inactivation of *Ythdc2*

The *Ythdc2* targeting construct was designed to insert a *loxP*-flanked hygromycin phosphotransferase-thymidine kinase (HyTK) drug selection cassette into *Ythdc2* intron 12, and a *loxP* site into intron 16 (Figure S1A). Genomic fragments were amplified from the *Ythdc2*-containing BAC clone RP23-138L17 by PCR with high-fidelity Taq DNA polymerase (BD Biosciences). The targeting construct was confirmed by sequencing. *FspI*-linearized targeting construct was electroporated into V6.5 mouse embryonic stem (ES) cells, and ES cells were cultured in media containing 120 µg/ml hygromycin B (Invitrogen). Of 92 hygromycin-resistant ES clones screened by long-range PCR, one clone (C4) was homologously targeted. Clone C4 was expanded and electroporated with *Cre-*expressing plasmid pOG231, followed by culture in media containing 2 µM ganciclovir. Ninety-two clones were screened for removal of the HyTK cassette and presence of *loxP* sites flanking *Ythdc2* exons 13-16, resulting in nine positive clones (*Ythdc2*^fl/+^). Two (C4B9 and C4H7) *Ythdc2*^fl/+^ ES clones were injected into blastocysts. The resulting chimeric mice transmitted the *Ythdc2* floxed allele through the germline.

*Ythdc2* floxed mice were crossed with four different Cre strains: *Actb*-Cre (Lewandoski et al., 1997), *Stra8*-Cre (Lin et al., 2017), *Neurog3*-Cre (Schonhoff et al., 2004), and *Ddx4*-Cre^ERT2^ (John et al., 2008). *Ddx4-*Cre^ERT2^ is tamoxifen-inducible Cre and expressed throughout spermatogenesis, including spermatogonia, spermatocytes, and round spermatids (John et al., 2008). Tamoxifen injection induces Cre-mediated deletion of exons 13-16 (Figure S1A). As described previously (Guan et al., 2020; Shi et al., 2019), tamoxifen (Sigma) was suspended with corn oil (Sigma) and injected intraperitoneally daily into the ≥8-week-old *Ythdc2*^fl/-^ *Ddx4-*Cre^ERT2^ males (2 mg/30 g body weight) for five consecutive days. *Ythdc2*^fl/-^ males (no Cre) injected with tamoxifen were used as controls. Testes were collected for analysis at 2, 4, 6, 8, and 10 days post-tamoxifen treatment. *Ythdc2* wild type (473 bp) and floxed (396 bp) alleles were assayed with primers GATGGGCCATATTCCACTTG and TGCTGCTGTTGGCTTTTATG. *Ythdc2*^-^ (deletion) allele (426 bp) was assayed with primers GCCAGTGTTCCAAAATTGCT and TGCTGCTGTTGGCTTTTATG. *Ddx4-*Cre^ERT2^ allele (205 bp) was assayed with primers ATACCGGAGATCATGCAAGC and GGCCAGGCTGTTCTTCTTAG.

### Histology, TUNEL, immunofluorescence, and nuclear spread analyses

For histological analysis, testes were fixed in Bouin’s solution at room temperature overnight, embedded in paraffin and sectioned in 6 μm. Sections were stained with hematoxylin and eosin (Fisher Scienctific). For TUNEL and immunofluorescence assays, testes were fixed in 4% paraformaldehyde (in 1× PBS) at 4°C overnight, dehydrated in 15% sucrose (in 1× PBS) for 3 hours, and finally in 30% sucrose (in 1× PBS) overnight. Dehydrated testes were embedded in Optimal Cutting Temperature compound (OCT) and cut in 6-μm sections on a cryostat. Alternatively, for TUNEL and immunofluorescence assays, testes were fixed in the modified Davidson’s fixative solution (30% of a 37-40% formaldehyde solution, 15% ethanol, and 5% glacial acetic acid) overnight, embedded in paraffin, and cut in 6-μm sections on a microtome. After deparaffinization and rehydration, slides were immersed in the antigen retrieval buffer (10 mM sodium citrate buffer, 0.05% Tween-20, pH 6.0) at 95°C for 20 minutes (Slides used for TUNEL assay were not processed by heat-induced antigen retrieval). For permeation treatment, slides were incubated in 0.5% Triton X-100 in 1× PBS at room temperature for 10 minutes. The slides were then blocked with 10% goat serum in PBST at 37°C for 1 hour followed by incubating with anti-SP10 antibodies at 37°C overnight. The morphology of acrosome revealed by anti-SP10 (ACRV1) antibody was used to precisely identify stages of seminiferous tubules. Slides were rinsed with PBS containing 0.1% tween-20 and then incubated with goat anti-rabbit, goat anti-mouse, or goat anti-guinea pig secondary antibodies (Vector Lab) at 37°C for 1h. Mounting medium with DAPI was added to the slides for imaging. For surface nuclear spread analysis, spermatocytes from 2, 4 and 6 dpt testes as well as the control were used according to the method described elsewhere (Dia et al., 2017; Peters et al., 1997) with minor modifications. Briefly, testicular tubules were separated from testis after removing the tunica albuginea and soaked in nuclear extraction hypotonic buffer (30 mM Tris, 50 mM Sucrose, 17 mM Trisodium Citrate Dihydrate, 5 mM EDTA, 0.5 mM dithiothreitol (DTT), 1 mM phenylmethylsulfonyl fluoride (PMSF)). Cells were released by grinding with tweezers and suspended in 100 mM sucrose, and were then spread on a thin layer of paraformaldehyde solution containing Triton X-100 on slides. The primary antibodies used for immunofluorescence analyses are listed in Key Resources Table.

### Imaging analysis

Histological images were captured on the Leica DM5500B microscope with a DFC450 digital camera (Leica Microsystems). Most immunolabeled chromosome spread images were taken on a Leica DM5500B microscope with an ORCA Flash4.0 digital monochrome camera (Hamamatsu Photonics). Images were processed using Adobe Photoshop CS6 and ImageJ v1.52p software packages (NIH, USA). Confocal images were acquired on a Leica SP5 confocal system with a 100x (1.46 NA) objective lens and processed with Huygens Essential deconvolution software (Scientific Volume Imaging). For supplemental movies, surface 3D renderings were created with Imaris software (Bitplane) and rotation sequences exported.

### YTHDC2 antibody production

Recombinant 6×His-YTHDC2 fusion protein (residues 1 to 239; mouse YTHDC2 protein accession number, NP_001156485.1) was expressed in *E. coli* BL21 cells using the pQE30 vector. The cDNA fragment encoding the N-terminal 239 aa of mouse YTHDC2 was amplified by PCR from the *Ythdc2* plasmid (NM_001163013) (Cat. No. MR226474, OriGene Technologies) using these two primers: 5’-CGCGGATCCATGTCCAGGCCGAGCAG-3’ and 5’-CCCAAGCTTTGTGGTCTTTCCAGACCCAGT-3’, and was cloned into the *Bam*HI and *Hind*III sites of the pQE30 vector. The purified 6×His-YTHDC2 recombinant protein was used to immunize two rabbits, yielding antisera UP2562 and UP2563 (Cocalico Biologicals Inc.). Affinity-purified antiserum UP2563 was used for immunofluorescence and Western blotting analyses in this study.

### Purification of pachytene spermatocytes by STA-PUT

Pachytene spermatocytes were purified from control (*Ythdc2*^fl/-^; heterozygote) and *Ythdc2*^iKO^ adult testes at 2 dpt using a mini-STA-PUT method (Bellve, 1993; Bryant et al., 2013; La Salle et al., 2009). Each purification was performed on testes from two or three adult males. Triplicate purifications were carried out per genotype. The purified spermatocytes were examined for purity by immunofluorescence of SYCP3 and γH2AX, and DAPI staining on surface nuclear surface spreads. The spermatocyte samples with a purity of more than 80% pachytene cells were used for RNA-seq and Western blot analyses.

### RNA-Seq library construction and sequencing

Total RNA of the purified pachytene spermatocytes was extracted using Trizol reagent (Ambion) according to the manufacturer’s instructions. The RNA quality and quantity were assessed using the Agilent 2100 Bioanalyzer. The cDNA library was constructed with 3 μg of input total RNA for each sample using the Illumina TruSeq RNA Sample Preparation Kit according to the manufacturer’s instructions. The peak and average final fragment sizes were ∼280 bp and ∼400 bp, respectively. 100-cycle sequencing of the libraires was performed on Illumina NovaSeq 6000 (Illumina, San Diego, USA) at the Next-Generation Sequencing Core (NGSC) at the University of Pennsylvania.

### Bioinformatic analysis

Reads were mapped to the mouse genome (GRCm38, Ensembl release 95) using salmon v1.3.0 (salmon quant with options -l A –validateMappings–gcBias) (Patro et al., 2017). Further analysis was performed using R (https://www.r-project.org/) version 3.6.3 and Bioconductor (https://bioconductor.org/). The DESeq function of DESeq2_1.25.10 bioconductor package (Love et al., 2014) was used to obtain log2 fold changes of gene expression between control (*Ythdc2*^fl/-^) and mutant (*Ythdc2*^iKO^)samples and the adjusted *P* values. Adjusted *P* value 0.1 was used as a threshold for statistical significance. Only protein-coding genes with average expression >= 50 across samples were analyzed. In MA plots the genes with significantly different expression between the *Ythdc2*^iKO^ and control samples with log2 fold change >= 0.5 or <= (−0.5) were highlighted. The volcano plots were plotted using EnhancedVolcano function from EnhancedVolcano 1.3.5 package (https://github.com/kevinblighe/EnhancedVolcano). Significantly differentially expressed genes with the log2 fold change >= 0.5 or <= (−0.5) in the *Ythdc2*^iKO^ were highlighted. The group of genes found to be significantly up- or downregulated in the *Ythdc2*^iKO^ were searched for enriched Gene Ontology terms using ENRICHR (Chen et al., 2013; Kuleshov et al., 2016) and the enriched categories were shown. To find out whether the dysregulated genes in the *Ythdc2*^iKO^, are expressed in the specific spermatogenic populations, we re-analyzed the published dataset containing gene expression data for spermatogonial stem cells, spermatocytes and round spermatids (ENA: PRJEB15333) (Davis et al., 2017) and plotted the expression of *Ythdc2*^iKO-^ dysregulated genes in these populations. To see whether the *Ythdc2*^iKO^ dysregulation correlates with the transcript m^6^A levels, we compared the *Ythdc2*^iKO^ expression changes between the genes reported to be enriched or depleted of m^6^A (GSE102346) (Wojtas et al., 2017). RNA-seq data of pachytene spermatocytes were deposited to GEO under GSE166568.

### qRT-PCR analysis

Testes and pachytene spermatocytes from control (*Ythdc2*^fl/-^) and *Ythdc2*^iKO^ (*Ythdc2*^f/-^ *Ddx4*-Cre^ERT2^) adult mice at 2 dpt were used for total RNA extraction. 1 μg of total RNA from each sample was treated with RNase-free DNase I (Thermo Fisher Scientific Inc., USA), used for reverse transcription using the RevertAid First Strand cDNA Synthesis Kit (Thermo Fisher Scientific Inc., USA) with oligo(dT)_18_ primers in a final volume of 20 μl, and diluted by addition of 80 μl H_2_O. The quantitative real-time reverse transcription PCR (qRT-PCR) was performed using the PowerUp^TM^ SYBR Green master mix (Applied Biosystems, Thermo Fisher Scientific Inc., USA) according to the manufacturer’s instructions in technical triplicate for each gene in an optical 384-well plate on the QuantStudio™ Real-Time PCR 6 Flex System (Applied Biosystems). Each real-time PCR reaction (in a final volume of 10 μl) contained 5 μl 2 × SYBR Green Master Mix, 0.25 μl of each primer (10 μM), 0.5 μl cDNA, and 4.0 μl H_2_O.

Primers were designed using the NCBI/Primer-BLAST with these parameters: PCR amplicon length of 100 - 250 bp, melting temperature (Tm) at approximately 60 °C, and primer pair specificity check against mouse mRNA Refseq database. The 2^−ΔΔCt^ method was used to calculate the relative expression of genes. Gene expression levels were normalized to the *Actb* level. The specific primers used for qRT-PCR are listed in Table S3.

### Western blot analysis

Adult testes or spermatocytes purified by STA-PUT were homogenized in 4 volumes of protein extraction buffer (62.5 mM Tris-HCl (pH 6.8), 3% SDS, 10% glycerol, 5% 2-mercaptoethanol). Samples were boiled at 95°C for 10-15 minutes and centrifuged at 12,000 g for 10 minutes. 30 μg of protein extracts per sample was resolved by SDS-PAGE and transferred onto nitrocellulose membranes using semi-dry transferring using iBlot system or onto polyvinylidene fluoride (PVDF) membranes by wet transferring in transfer buffer (25 mM Tris, 192 mM glycine, 0.02% SDS, 10% methanol, pH8.3). The primary antibodies used for western blotting analysis are listed in Key Resources Table.

### Culture of spermatocytes and inhibitor treatment

Freshly dissected testes from 2 to 3-month-old male mice were decapsulated and placed into Krebs buffer (120 mM NaCl, 4.8 mM KCl, 25.2 mM NaHCO_3_, 1.2 mM KH_2_PO_4_, 1.2 mM MgSO_4_, 1.3 mM CaCl_2_, and 11.1 mM dextrose) containing collagenase (0.5 mg/ml). Testicular tubules were digested at 32°C for about 10 min and were transferred into Krebs buffer containing trypsin (0.25 mg/ml) and DNase (0.02 mg/ml). Cells were incubated at 32°C for another 10 min with repeated pipetting to become single-cell suspension. The cells were passed through a 100-μm nylon mesh and washed twice in Kreb’s buffer before culture.

5×10^6^ adult mouse testicular cells were cultured in 1 ml MEMα (minimal essential media with alpha modifications) containing 5% streptomycin, 7.5% penicillin, 0.29% DL-lactic acid sodium salt, 0.59% HEPES, and 5% fetal bovine serum in one well of a six-well plate in 5% CO_2_ at 32°C. After an one-hour acclimation culture period, cells were then treated with inhibitors. All the treatments were performed three times.

For the metaphase I competency assay, testicular cells were treated with 4 μM okadaic acid (from a stock of 244 μM in ethanol). Control cultures received an equivalent volume of ethanol alone. Five hours after culture with okadaic acid, cells were harvested for nuclear spread analysis.

For the microtubule depolymerization assay, testicular cells were treated with 0.1 μg/ml nocodazole (from a stock of 0.1 mg/ml in DMSO). Control cultures received an equivalent volume of DMSO. 16 hours after nocodazole treatment in culture, cells were harvested for nuclear spread analysis.

For the actin depolymerization assay, cells were treated with 2.5 μg/ml cytochalasin D (from a stock of 2 mg/ml in DMSO) or 0.5 μM latrunculin A (from a stock of 0.5 mM in DMSO). Control cultures received an equivalent amount of DMSO. 16 hours after inhibitor treatment, cells were harvested for nuclear spread analysis.

### Statistical analysis

The data were expressed as the mean ± std and were statistically analyzed with Student’s *t*-test using SPSS statistics software version 26 (IBM Corp., USA). Differences with *P* < 0.05 were considered to be statistically significant.

## Supplementary information

Figure S1-S7

Table S1 to S3 (uploaded separately)

Movie S1 and S2 (uploaded separately)

## Supplementary Tables

**Table S1.** Differentially expressed genes in Ythdc2^iKO^ pachytene spermatocytes.

**Table S2.** Overlap of YTHDC2 RIP-seq target transcripts with RNA-seq differentially expressed genes in Ythdc2^iKO^ (2 dpt) pachytene spermatocytes.

**Table S3.** qRT-PCR primers

**Movie S1. Deconvoluted confocal image of a *Ythdc2*^fl/-^ pachytene spermatocyte. Related to Figure 2.**

The synaptonemal complex (SYCP3), red; Centromere (CREST), green; DNA (DAPI), blue. Note the random distribution of centromeric chromosomal ends.

**Movie S2. Deconvoluted confocal image of a pachytene spermatocyte from *Ythdc2*^iKO^ (4 dpt) testis. Related to** **Figure 2**.

The synaptonemal complex (SYCP3), red; Centromere (CREST), green; DNA (DAPI), blue. Note clustering of centromeric chromosomal ends to one side of the nucleus.

## Supplementary figures

**Figure S1.**
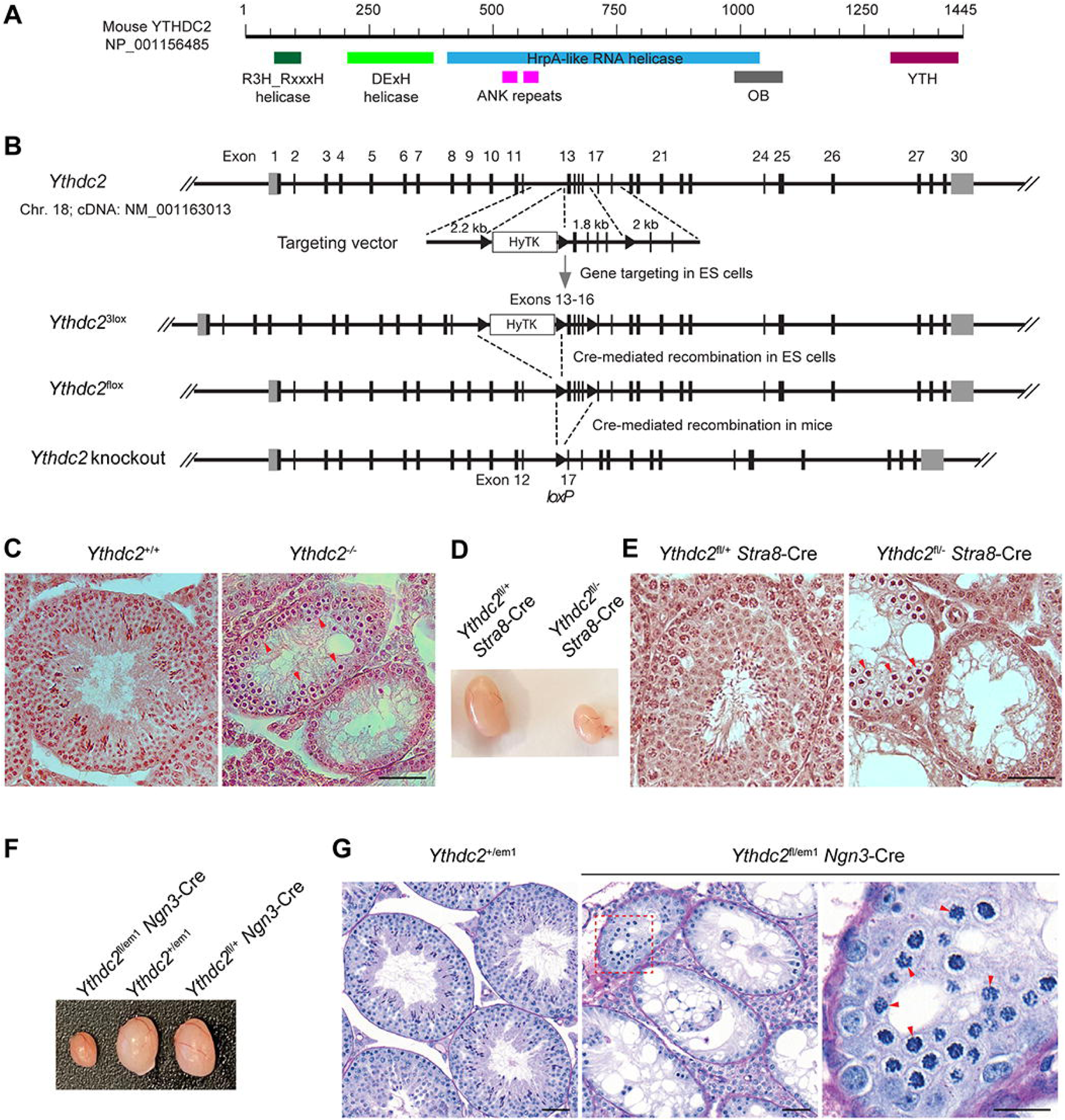
*Ythdc2* Conditional Knockout Strategy. Related to Figure 1. (A) Illustration of functional domains in the mouse YTHDC2 protein. (B) Schematic diagram of mouse *Ythdc2* gene structure, targeting vector, and targeted alleles. *Ythdc2* consists of 30 exons. Exons 13-16 are flanked by *loxP* sites. Deletion of exons 13-16 results in a deletion of 149 amino acids (aa 578 – 726) and a frameshift in the mutant transcript. Coding exons are shown as black bars. 5’ and 3’ untranslated regions are shown as gray boxes. HyTK is a double selection marker and enables hygromycin positive selection and thymidine kinase-negative selection in ES cells. (C) Histology (H&E staining) of 3-month-old wild-type and *Ythdc2*^-/-^ testes. Arrowheads indicate abnormal metaphase-like spermatocytes. (D) Reduced testis size in 6-week-old *Ythdc2*^fl/-^ *Stra8*-Cre males. (E) Histology (H&E staining) of 6-week-old control and *Ythdc2*^fl/-^ *Stra8*-Cre testes. Arrowheads indicate abnormal metaphase-like spermatocytes. (F) Reduced testis size in 8-week-old *Ythdc2*^fl/em1^ *Ngn3*-Cre mice. (G) Periodic acid Schiff-stained sections of Bouin’s-fixed testes from 5-month-old *Ythdc2*^fl/em1^ *Ngn3*-Cre and *Ythdc2*^+/em1^ mice. *Ythdc2*^em1^ is a deletion allele (Jain et al., 2018). In the higher magnification image of the boxed region, arrowheads annotate abnormal metaphase-like cells. Scale bars (C, E, and G left panels), 50 μm; (G right panel), 20μm.

**Figure S2.**
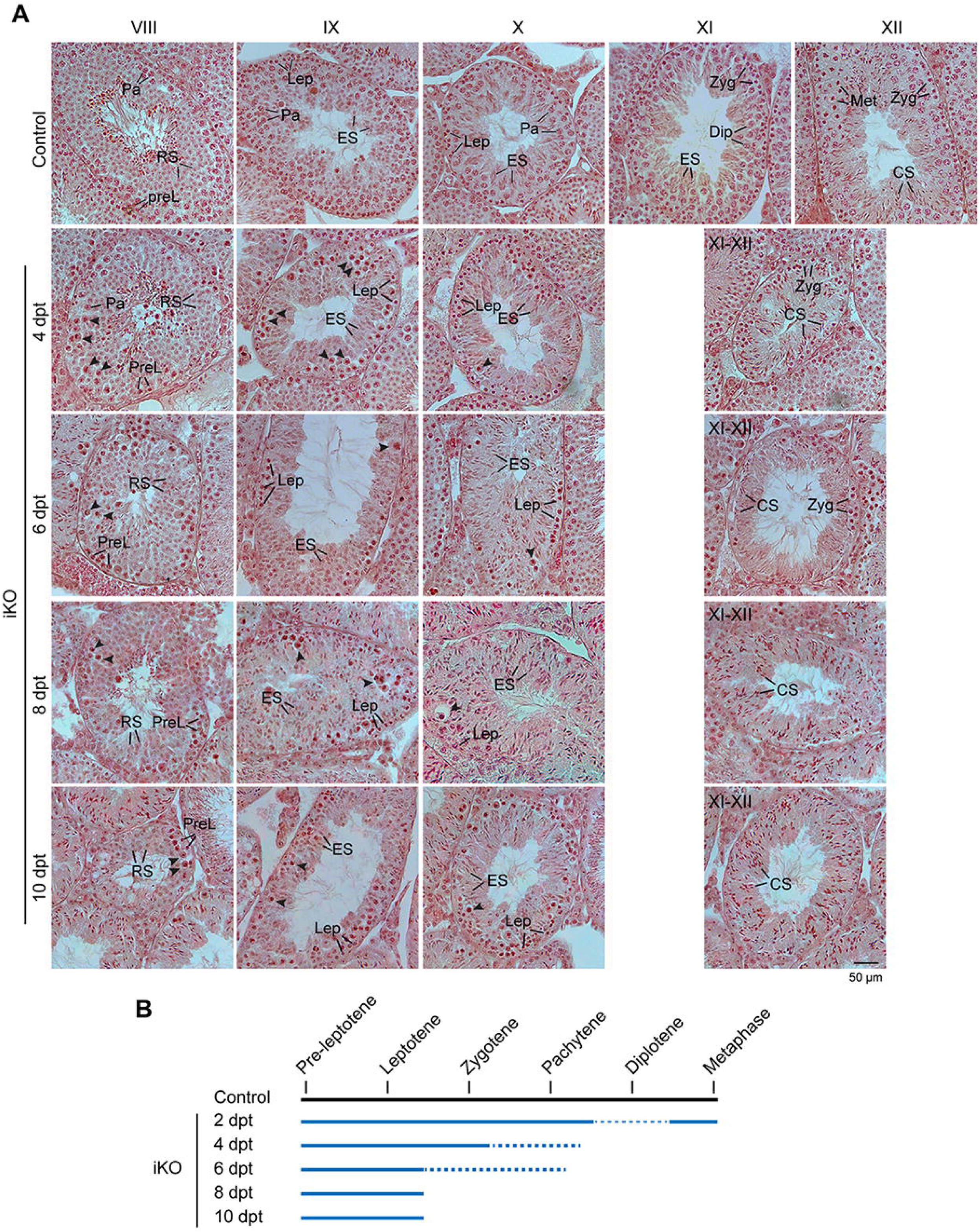
Histological Analysis of Control (*Ythdc2*^fl/-^, no Cre) and *Ythdc2*^iKO^ Adult Testes at 4, 6, 8, and 10 dpt. Related to Figure 1. (A) Histology of stage VIII, IX, X, XI, and XII tubules are shown. For 6, 8, and 10 dpt, stages XI and XII could not be distinguished from each other, because of a total loss of diplotene and metaphase spermatocytes. Note a partial loss of zygotene spermatocytes in stages XI/XII at 6 dpt and a complete loss of zygotene spermatocytes in stages XI/XII at 8 and 10 dpt. Arrowheads indicate the apoptotic spermatocytes. Abbreviations: PreL, preleptotene; Lep, leptotene; Zyg, zygotene; Pa, pachytene; Dip, diplotene; Met, metaphase I or II spermatocyte; RS, round spermatid; ES, elongating spermatid; CS, condensing spermatid. Scale bar, 50 μm. (B) Time-dependent progressive loss of spermatocytes in adult tamoxifen-treated *Ythdc2*^iKO^ testes. Lines indicate the presence of spermatocytes. Dash lines indicate reduction in the indicated type of spermatocytes. A thinner dash line at 2 dpt indicates a severe reduction in the number of diplotene spermatocytes.

**Figure S3.**
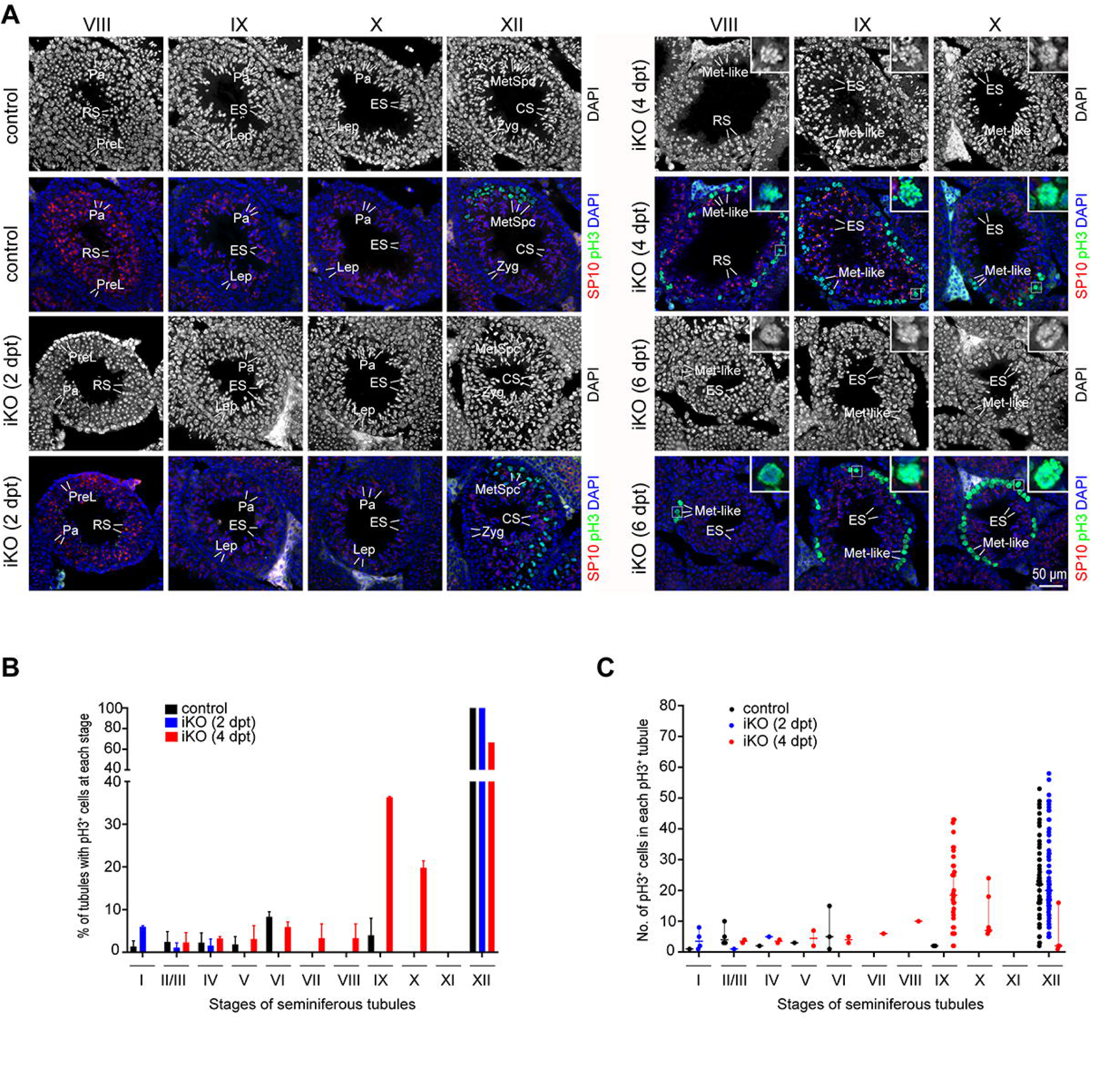
Abnormally Condensed Preleptotene/Leptotene Spermatocytes Are pH3-positive in Stage VIII-X Seminiferous Tubules from *Ythdc2*^iKO^ Testes at 4 dpt and 6 dpt. Related to Figures 3 and 7. (A) Immunofluorescence analysis of pH3 (green) in seminiferous tubules from adult control (*Ythdc2*^fl/-^) and *Ythdc2*^iKO^ testes at 2, 4, and 6 dpt. The acrosome (red) was stained with anti-SP10 antibody. Nuclear DNA was counterstained with DAPI (blue). Abbreviations: PreL, preleptotene; Lep, leptotene; Zyg, zygotene; Pa, pachytene; RS, round spermatid; ES, elongating spermatid; CS, condensed spermatid; Met, metaphase I or II spermatocytes; Met-like, metaphase-like spermatocyte. Inset shows an enlarged view of the boxed pH3-positive metaphase-like spermatocyte. Scale bar, 50 μm. (B) Percentage of pH3-positive tubules from adult control, 2 dpt *Ythdc2*^iKO^, and 4 dpt *Ythdc2*^iKO^ testes. The mean ± s.d. values were plotted. (C) The number of pH3-positive cells in pH3-positive tubules (mean ± s.d.) from adult control, 2 dpt *Ythdc2*^iKO^, and 4 dpt *Ythdc2*^iKO^ testes. (B, C) Two males per genotype were analyzed. At least 200 tubules were counted for each mouse. The stages of seminiferous tubules are shown in Roman numerals.

**Figure S4.**
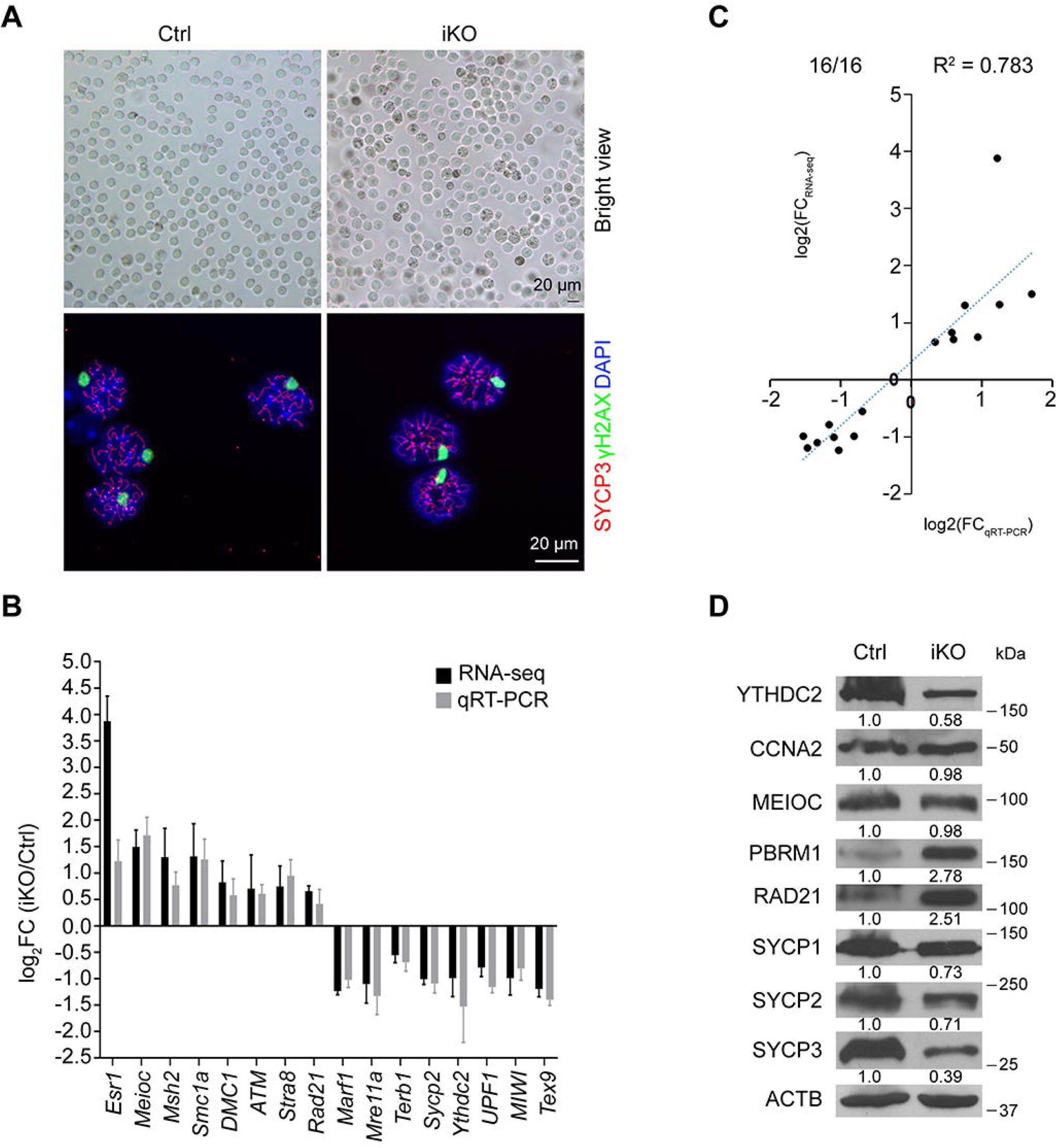
Validation of Differentially Expressed Genes in Purified Pachytene Cells by qRT-PCR and Western Blot Analyses. Related to Figure 5. (A) Assessment of purity of Sta-put isolated control (*Ythdc2*^fl/-^) and *Ythdc2*^iKO^ pachytene spermatocytes by bright view and spread analysis. (B) qRT-PCR analysis of a set of 8 upregulated genes and 8 downregulated genes in *Ythdc2*^iKO^ pachytene cells. (C) Correlation of expression changes of 8 upregulated and 8 downregulated genes between RNA-seq and qRT-PCR. (D) Western blot analysis of selected proteins in purified pachytene cells. Western blotting was performed on independent samples twice. ACTB serves as a loading control. Quantification of each protein is normalized to ACTB and set at 1.0 in control pachytene cells.

**Figure S5.**
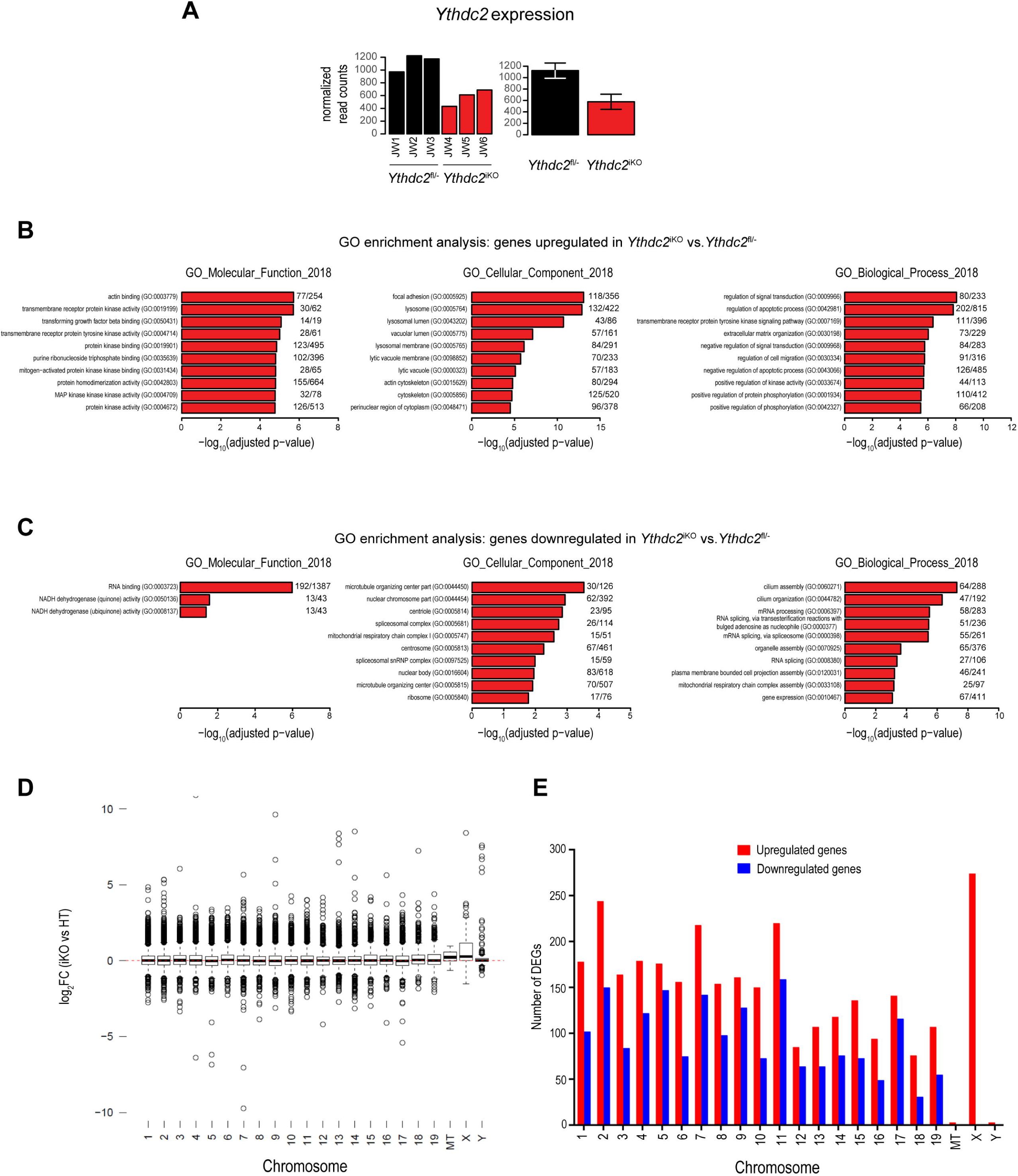
Gene Ontology Analysis of Differentially Expressed Genes in *Ythdc2*^iKO^ Pachytene Cells. Related to Figure 5. (A) Comparison of *Ythdc2* expression in individual samples. (B) Gene ontology analysis of the up-regulated genes in *Ythdc2*^iKO^ pachytene cells. (C) Gene ontology analysis of the down-regulated genes in *Ythdc2*^iKO^ pachytene cells. (D) Expression changes based on individual chromosomes. Mitochondria are included. (E) The number of upregulated vs downregulated genes on each chromosome and mitochondria.

**Figure S6.**
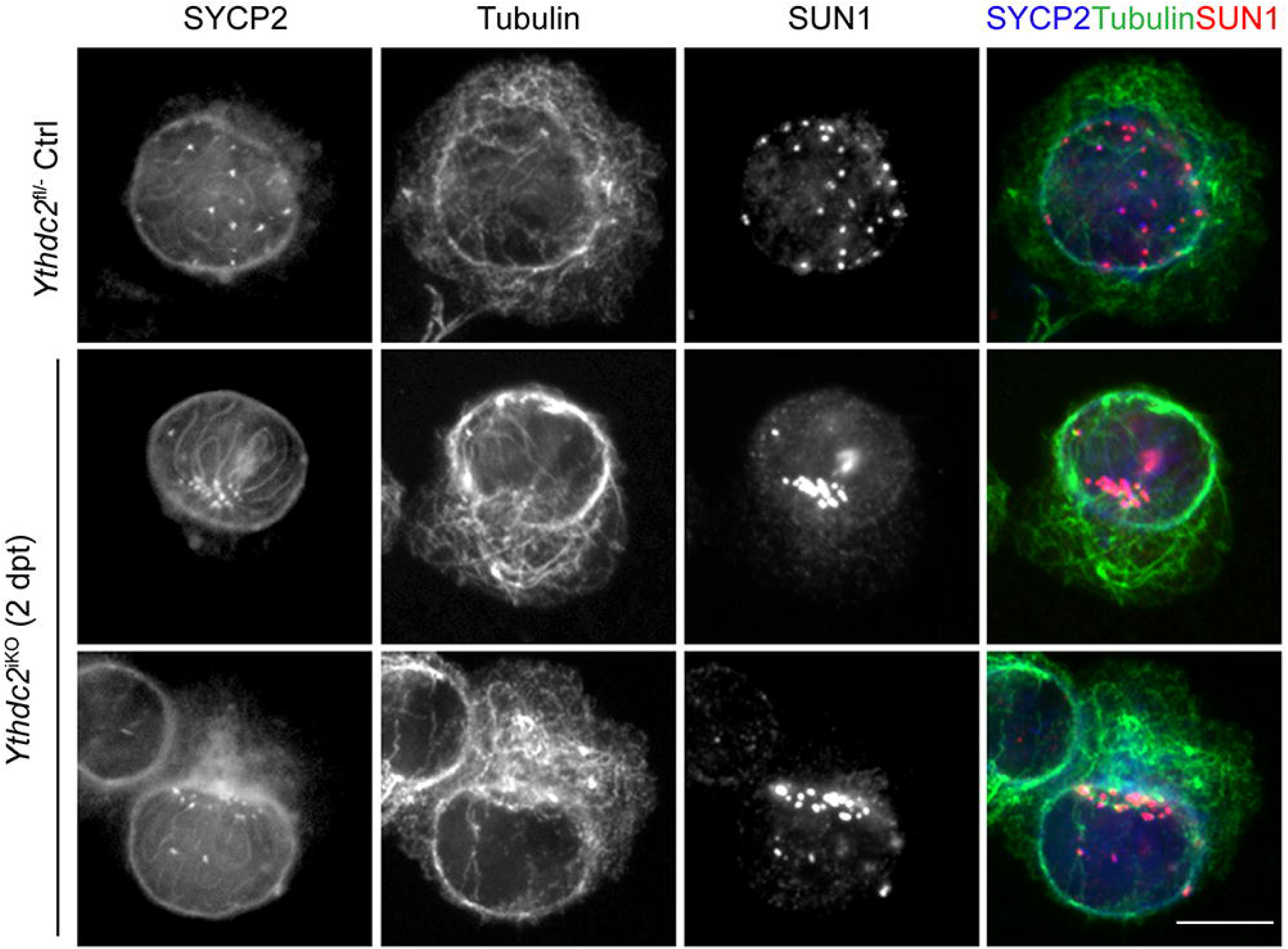
Polarized distribution of telomeres, microtubules, and cytoplasm in *Ythdc2*^iKO^ pachytene cells. Related to Figure 6. Immunofluorescence of α-tubulin, SUN1, SYCP2 was performed in intact control (*Ythdc2*^fl/-^) and *Ythdc2*^iKO^ (2 dpt) pachytene spermatocytes. Scale bar, 10 μm.

**Figure S7.**
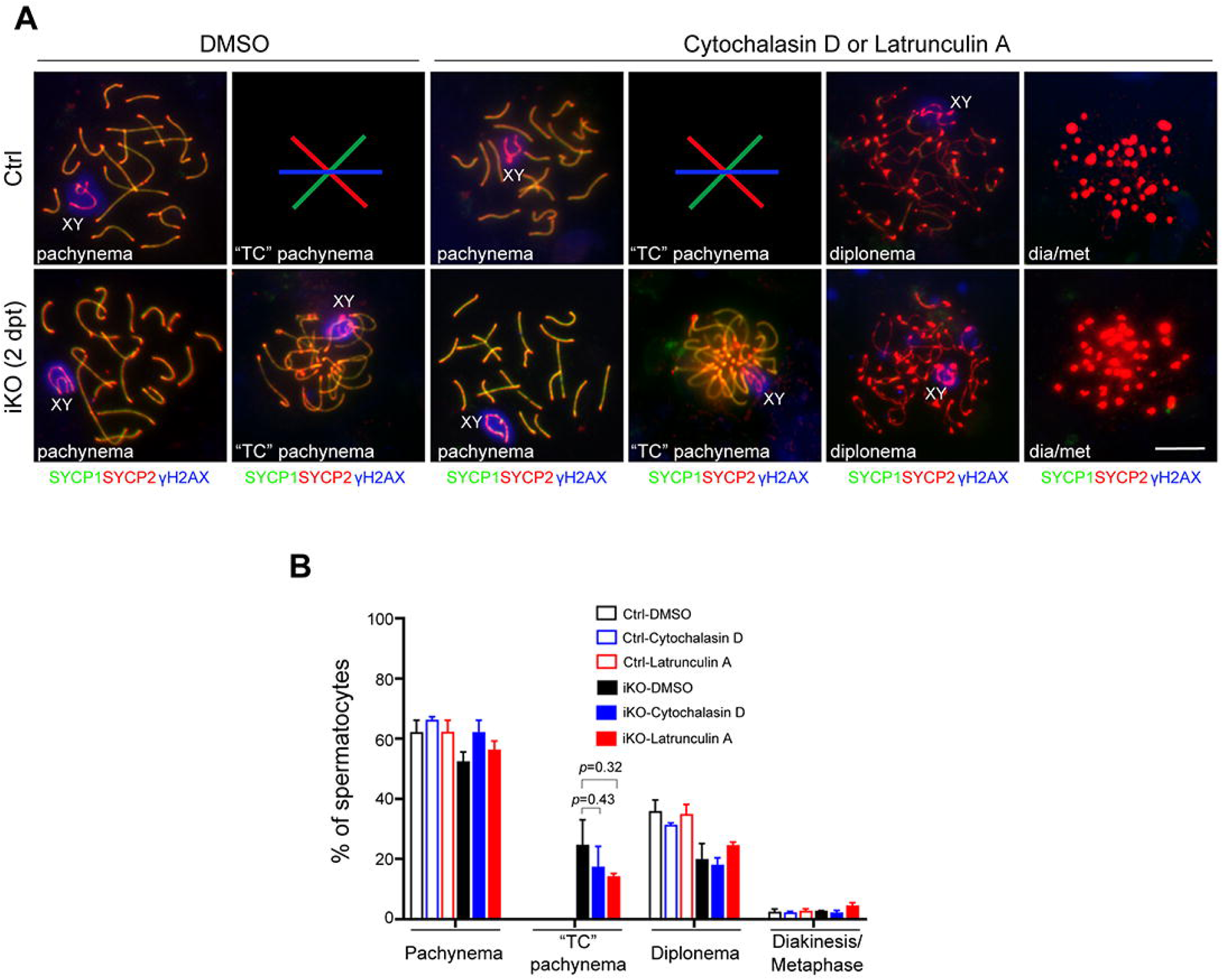
Treatment with inhibitors of actin filaments cytochalasin D or latrunculin A. Related to Figure 6. (A) Nuclear spread analysis of pachytene, diplotene, and diakinesis/metaphase I (dia/met) spermatocytes from control (*Ythdc2*^fl/-^) and *Ythdc2*^iKO^ (2 dpt) testes after treatment with DMSO, cytochalasin D, or latrunculin A. TC, telomere clustering. Scale bar, 10 μm. (B) Percentage of spermatocytes (mean ± s.d.). Only pachytene, diplotene, and diakinesis/metaphase I spermatocytes are included. At least 350 cells were counted per genotype per treatment group. Two males per genotype per experiment were used. The experiments were performed twice.

